# Integration of Protein Interactome Networks with Congenital Heart Disease Variants Reveals Candidate Disease Genes

**DOI:** 10.1101/2021.01.05.423837

**Authors:** Barbara Gonzalez-Teran, Maureen Pittman, Franco Felix, Desmond Richmond-Buccola, Reuben Thomas, Krishna Choudhary, Elisabetta Moroni, Giorgio Colombo, Michael Alexanian, Bonnie Cole, Kaitlen Samse-Knapp, Michael McGregor, Casey A. Gifford, Ruth Huttenhain, Bruce D. Gelb, Bruce R. Conklin, Brian L. Black, Benoit G. Bruneau, Nevan J. Krogan, Katherine S. Pollard, Deepak Srivastava

## Abstract

Congenital heart disease (CHD) is present in 1% of live births, yet identification of causal mutations remains a challenge despite large-scale genomic sequencing efforts. We hypothesized that genetic determinants for CHDs may lie in protein interactomes of GATA4 and TBX5, two transcription factors that cause CHDs. Defining their interactomes in human cardiac progenitors via affinity purification-mass spectrometry and integrating results with genetic data from the *Pediatric Cardiac Genomic Consortium* revealed an enrichment of *de novo* variants among proteins that interact with GATA4 or TBX5. A consolidative score that prioritized interactome members based on variant, gene, and proband features identified likely CHD-causing genes, including the epigenetic reader GLYR1. GLYR1 and GATA4 widely co-occupied cardiac developmental genes, resulting in co-activation, and the GLYR1 missense variant associated with CHD disrupted interaction with GATA4. This integrative proteomic and genetic approach provides a framework for prioritizing and interrogating the contribution of genetic variants in disease.

## INTRODUCTION

Birth defects are complex developmental phenotypes affecting about 6% of births worldwide, but their genetic roots are multifarious and difficult to ascertain (Christianson and Howson, 2006; Deciphering Developmental Disorders Study, 2015). Particularly challenging are rare disorders and more common but complex defects with high allelic and locus heterogeneity. In recent years, whole-exome sequencing has accelerated our understanding of such disorders, including the most common birth defect, congenital heart disease (CHD) (Zaidi et al., 2013; Heyne et al., 2018; Homsy et al., 2015; Jin et al., 2017; Richter et al., 2020). *De novo* monogenic aberrations were found to collectively contribute to ∼10% of CHD cases, whereas rare inherited and copy number variants have been identified in ∼1% and 25% of cases, respectively (Zaidi and Brueckner, 2017). Additionally, polygenic and oligogenic inheritance models where multiple genetic variants with epistatic relationships are implicated, have been proposed as mechanistic explanations for certain complex phenotypes. A recent study from our group highlighted the involvement of genetic modifiers in human cardiac disease (Gifford et al., 2019), but the net contribution of oligogenic inheritance remains to be determined. Although the growing catalogue of human genome variants has led to significant advances in our understanding of the genetic underpinnings of CHD, the cause of over 50% of CHD cases remains unknown (Zaidi and Brueckner, 2017).

A barrier to a complete understanding of CHD’s etiology is its immense genetic heterogeneity. Estimates based on *de novo* mutations alone indicate that more than 390 genes may contribute to CHD pathogenesis (Homsy et al., 2015). This heterogeneity reduces the statistical power of CHD risk gene analysis with the cohorts currently available. Recent work suggests that cohorts of approximately 10,000 parent-proband trios would be needed for whole-exome sequencing to detect ∼80% of genes contributing to haplo-insufficient syndromic CHD (Sifrim et al., 2016), highlighting the need for alternative strategies to identify CHD risk genes and to prioritize for potentially causative variants.

Many diseases display tissue-restricted phenotypes, but are rarely explained by mutations in genes with tissue-specific expression (Hekselman and Yeger-Lotem, 2020). Many cardiac malformations have been linked to variants in tissue-enriched cardiac transcription factors (cTFs) that are expressed more widely. Such cTFs typically form complexes with other tissue-enriched and ubiquitous proteins to orchestrate specific developmental gene programs (Lambert et al., 2018). cTF missense variants may disrupt specific interactions with other proteins, affecting their transcriptional cooperativity and causing disease (Ang et al., 2016; Garg et al., 2003; Moskowitz et al., 2011; Waldron et al., 2016). This observation suggests a functional relevance for cTF-interactors in genetic disorders, including CHD. Accordingly, an excess of protein-altering *de novo* mutations from the Pediatric Cardiac Genomic Consortium’s CHD cohort were found in ubiquitously expressed chromatin regulators that partner with cTFs to regulate the expression of key developmental genes (Zaidi et al., 2013). This led us to hypothesize that protein-protein interactors of cTFs associated with CHD may be enriched in disease-associated proteins, even if these proteins are not tissue-specific.

GATA4 and TBX5 are two essential cTFs (Kuo et al., 1997; Bruneau et al., 1999, 2001; Molkentin et al., 1997; Mori et al., 2006) and among the first monogenic etiologies of familial CHD (Li et al., 1997; Basson et al., 1997; Garg et al., 2003). Pathogenic variation in *TBX5* are a cause of septation defects and other forms of CHD in the setting of Holt-Oram syndrome (Basson et al., 1997; Li et al., 1997). Pathogenic variation in *GATA4* also causes atrial and ventricular septal defects, as well as pulmonary stenosis and outflow tract abnormalities (Garg et al., 2003; Rajagopal et al., 2007; Tomita-Mitchell et al., 2007). Subsequent studies have demonstrated that TBX5 and GATA4 cooperatively interact on DNA throughout the genome to regulate heart development (Garg et al., 2003; Ang et al., 2016). Disruption of the physical interaction between these cTFs or with other specific co-factors by missense variants can impair transcriptional cooperativity and lineage specification, and ultimately cause cardiac malformations (Ang et al., 2016; Garg et al., 2003; Maitra et al., 2009; Waldron et al., 2016). Therefore, the identification of human GATA4 and TBX5 (GT) protein interactors during cardiogenesis could highlight disease mechanisms and provide a powerful filter for interrogating the impact of protein-coding variants in CHD etiologies.

Here, we leveraged an integrated proteomics and human genetics approach that dissects the protein-protein interactors of endogenous GATA4 and TBX5 in human cardiac progenitor cells to identify and prioritize potential disease genes harboring CHD-associated variants, revealing novel aspects of cardiac gene regulation. This approach can be leveraged to study the genetic underpinnings of many human diseases.

## RESULTS

### Identification of the GATA4 and TBX5 Protein Interactomes in Cardiac Progenitors

Although many protein partners of GATA4 and TBX5 have been found in mice, the protein interactors that titrate their effects in early human cardiogenesis have yet to be systematically explored. To fill this gap in knowledge, we identified the GATA4 and TBX5 protein interactome (GT-PPI) in human induced pluripotent stem cell–derived cardiac progenitors (CPs) using antibodies against each endogenous cTF for affinity purification and mass spectrometry (Figure 1A). Using CRISPR Cas9-gRNA ribonucleoproteins, we generated clonal *TBX5* or *GATA4* homozygous knockout (KO) hiPSC lines as negative controls. These control lines were differentiated to CP and cardiomyocyte (CM) stages, and the absence of the respective cTF expression was confirmed (Figure S1A–E). Consistent with previous reports using murine cells (Luna-Zurita et al., 2016; Narita et al., 1997), *GATA4* and *TBX5* KO cells were able to differentiate into CMs, albeit with delayed beating and reduced differentiation efficiency (Figure S1E–G and Table S1 and S2).

**Figure 1:**
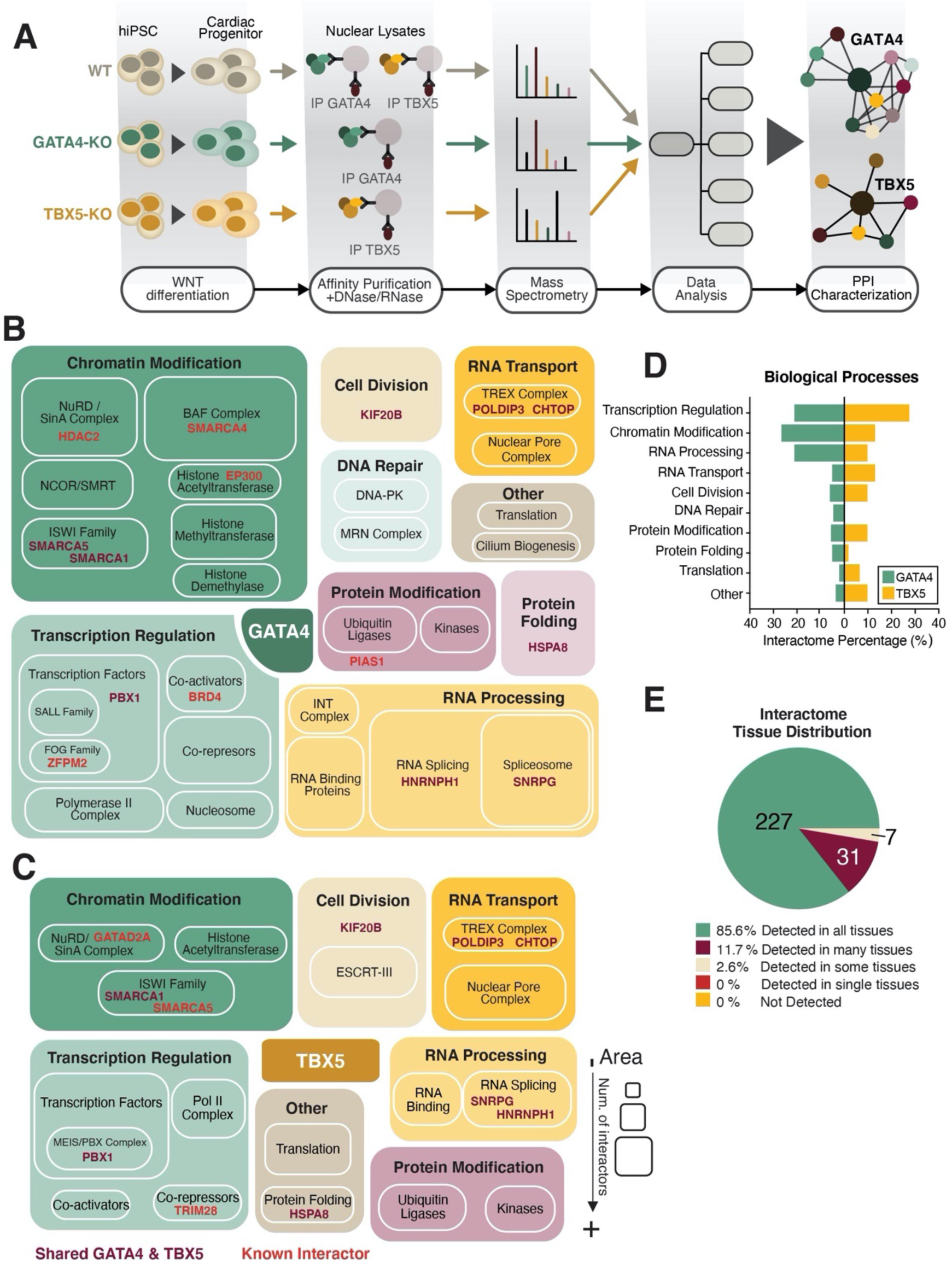
Generation of GATA4 and TBX5 protein interactomes in human iPSC-derived cardiac progenitors. (A) WTC11 and WTC11 CRISPR/Cas9 engineered clonal hiPSC lines (GATA4-KO & TBX5-KO) were differentiated to cardiac progenitors (CPs, differentiation day 6). CPs from differentiations that passed QC thresholds (see Methods) were subjected to affinity purification (AP) of endogenous GATA4 or TBX5 and their protein complexes from nuclear lysates treated with Benzonase (DNase/RNase enzyme). For each AP condition, replicates from three independent differentiations were analyzed by mass spectrometry (LC/MS). Affinity purification-mass spectrometry (AP-MS) results from KO CPs served as negative controls to remove antibody-specific background from the experimental samples’ signal; data were subjected to several further filtering steps to identify high-confidence GATA4 and TBX5 protein-protein interactome networks (PPINs). (B) GATA4 and (C) TBX5 interactors were manually annotated for biological processes and protein complexes based on literature available. Boxed areas are roughly proportional to the number of interactors they represent. Proteins interacting with both GATA4 and TBX5 (purple) and previously reported interactors (red) are highlighted. (D) Distribution of GATA4 and TBX5 PPIs across biological processes, as annotated in panels B & C. (E) Tissue expression distribution of GATA4 and TBX5 interactors across the six Human Protein Atlas categories based on transcript detection (NX≥1) in all 37 analyzed tissues (See Methods). Detected in single: detected in a single tissue; Detected in some: detected in more than one but less than one third of tissues; Detected in many: detected in at least a third but not all tissues; Detected in all: detected in all tissues; Not detected.

GATA4 or TBX5 mass spectrometry data were generated from three replicates of nuclei-enriched Day 6 hiPSC-derived cardiac progenitor wild-type (WT) or KO samples treated with RNase and DNase to focus on nucleic acid independent interactions (Figure 1A). An initial list of GT-interactors in WT CPs was obtained by scoring the proteins identified in WT affinity purification-mass spectrometry experiments to their corresponding KO control line using the established protein-protein interaction algorithm SAINTq (Teo et al., 2016). For further stringency, additional filtering was applied for the high-scoring interactors determined by SAINTq based on nuclear localization, co-expression in the same cells as the bait protein, and their differences in expression comparing WT and KO cells (See Methods). This approach yielded 252 proteins in total, which comprised several of the previously reported GATA4 and TBX5 interactors as well as novel interactors (Enane et al., 2017; Padmanabhan et al., 2020; Waldron et al., 2016). Mutations in several of these interactors have been already associated to human or mouse cardiac malformations, highlighting the potential of our approach for disease-gene discovery (Jin et al., 2017; Bouman et al., 2017; Castillo-Robles et al., 2018; Chen et al., 2020; Diets et al., 2019; Dsouza et al., 2019; Ferrante et al., 2006; Gordillo et al., 1993; Hinton et al., 2014; Homsy et al., 2015; Ji et al., 2020; Jones et al., 2012; Lebrun et al., 2018; Lei et al., 2012; Lepore et al., 2006; Maitra et al., 2010; Parisot et al., 2010; Pierpont et al., 2018; Takeuchi et al., 2011; Thienpont et al., 2010; Van Dijck et al., 2019; Wilczewski et al., 2018) (Figure 1B and 1C, Figure S2A and S2B and Table S3A and S3B).

Consistent with the interdependence of GATA4 and TBX5 during cardiac development, their networks showed some overlap, but the bulk of the detected interactors were unique to each cTF (Figure S2C). Both networks were enriched in proteins involved in similar biological processes (Figure 1B–D and Figure S2A and S2B). The top two most represented processes were transcription regulation and chromatin modification (Figure 1D), as expected from GATA4 and TBX5’s well-established functions in gene regulation. Transcription regulators (∼20% of GATA4 interactors and ∼30% of TBX5 interactors) comprised TFs, co-activators, co-repressors and polymerase II complex-associated proteins. Both known and previously unreported low-abundance TFs were found to interact with GATA4 and/or TBX5, demonstrating the relatively high sensitivity of the affinity purification-mass spectrometry approach (Figure 1B and 1C, Figure S2A and S2B). Chromatin modifiers (∼25% or 15% of GATA4 or TBX5 interactors, respectively) predominantly belonged to ATP-dependent complexes, including SWI/SNF, NuRD-CHD and ISWI subfamilies. In addition, we found several histone-modifying enzymes in the GATA4-PPI, some of which had been previously described as interactors in other cell types (Enane et al., 2017) (Figure 1B and 1C, Figure S2A and S2B). The GT-PPIs mostly included proteins expressed ubiquitously, with a small number of tissue-enriched interactors (Figure 1E).

We identified two biological processes highly enriched in GT-PPIs and not previously associated with these TFs. For instance, a substantial number of RNA processing and splicing proteins interacted with GATA4. Unexpectedly, several proteins involved in nucleocytoplasmic RNA transport interacted with both TFs, including proteins from the nuclear pore complex (NPC) (Figure 1B and 1C, Figure S2A and S2B). The significance of these interactions will require further study.

Overall, our findings demonstrate the power of affinity purification-mass spectrometry to identify endogenous interactors of GATA4 and TBX5 in an unbiased fashion and support a model where cardiac-specific functions result from interactions between cardiac-enriched TFs and more ubiquitously expressed proteins.

### GATA4:TBX5-Protein Interactome Is Enriched in Proteins Harboring Likely Deleterious CHD-Associated *De Novo* Variants

To determine whether the GT-interactors derived from CPs might help predict genetic risk factors for CHD, we assessed their intersection with *de novo* variants (DNV) and very rare (MAF ≤ 10^-5^) inherited variants found in published CHD and control cohorts from the Pediatric Cardiac Genomic Consortium. This whole-exome sequencing database represents the largest available cohort of parent-offspring trios with over 2500 CHD probands and their parents (Jin et al., 2017). We used a permutation-based statistical test to analyze the frequency of mutations in GT-interacting proteins among the CHD probands compared to the control group (see Methods) (Figure 2A). The analysis indicated that the GT-interactors were significantly more likely to map to protein-altering DNVs found in the CHD cohort than in the control cohort (adjusted odds ratio (OR) G-PPI: 5.83; T-PPI: 3.42). By contrast, very rare inherited variants occurred in GT-PPIN proteins with the same frequency in control and CHD groups (adjusted OR G-PPI: 1.19; T-PPI: 0.96) (Figure 2B and Table S4).

**Figure 2:**
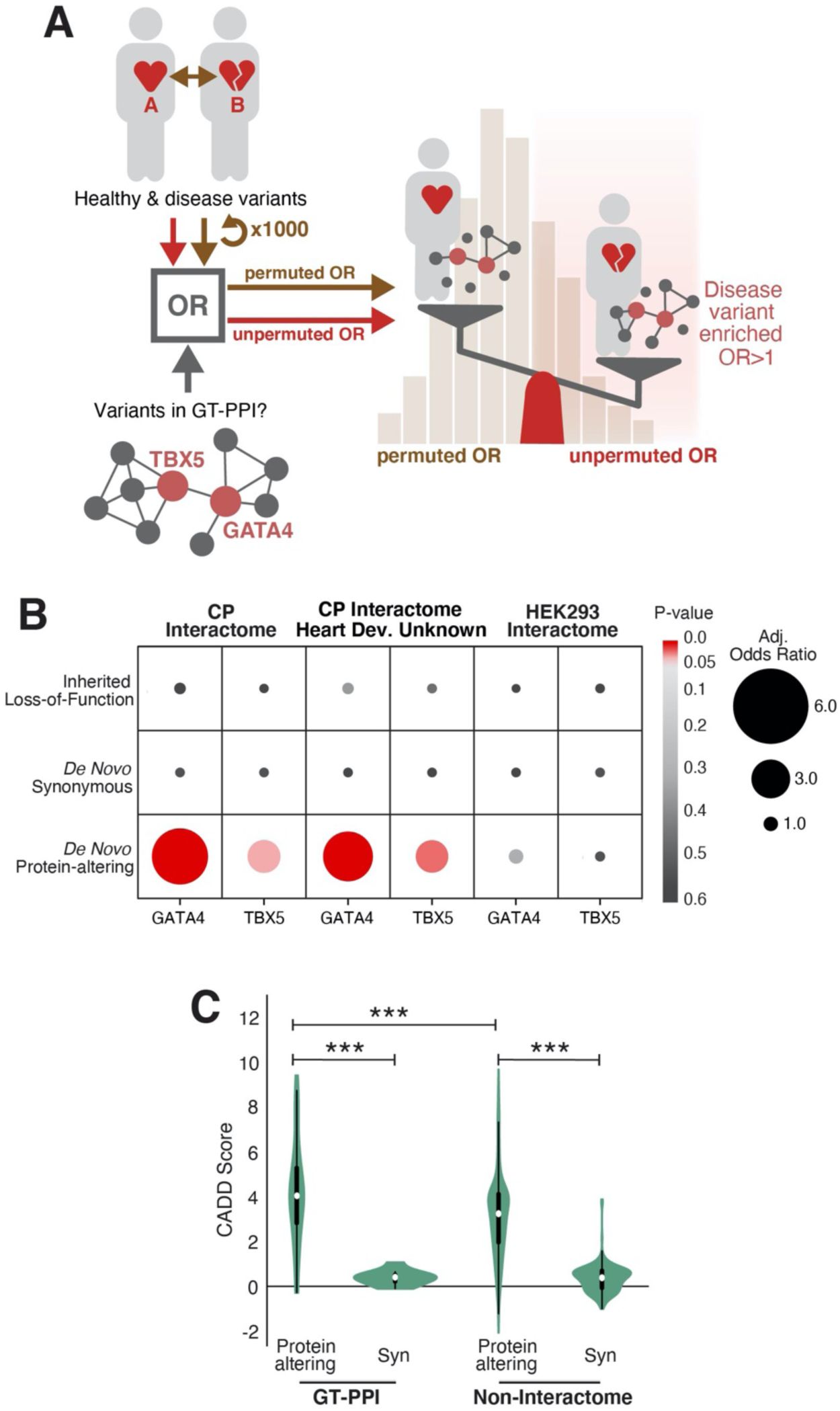
Enrichment of *de novo* variants among GATA4 and TBX5 interactome proteins among CHD trios. (A) Design of a permutation-based statistical test to analyze the enrichment in genetic variants from a CHD cohort relative to a control cohort in GATA4 or TBX5 PPIs (Odds Ratio, OR). The probability of CHD-associated genomic variation to occur by chance within the PPIs was calculated by randomly permuting the Pediatric Cardiac Genomic Consortium (PCGC) healthy (control group, A) and diseased (CHD group, B) IDs and calculating the corresponding ORs 1000 times. The enrichment in CHD-associated variants was then analyzed by calculating the unpermuted adjusted (Adj.) OR and its p-value. The Adj. OR is corrected for differences in sequencing depth across case and control datasets based on synonymous DNV counts (see Methods). (B) Permutation-based statistical test for different types of reported genomic variation (Inherited LoF, *de novo* synonymous and *de novo* protein-altering (non-synonymous)) from *PCGC* CHD and Control cohorts to analyze their enrichment within defined GATA4 and TBX5 PPIs. CP Interactome: group of proteins found as GATA4 or TBX5 interactors in CPs; CP Interactome Heart Dev. Unknown: CP Interactome after removing proteins included in a curated list of human/mouse cardiac development and CHD genes reported by the *PCGC* (Jin et al., 2017) (See Table S5); HEK293 Interactome: group of proteins found as GATA4 or TBX5 interactors in HEK293Ts. (C) Violin Plot representing the Combined Annotation-Dependent Depletion (CADD) scores for Protein-altering and Synonymous (Syn) variants found in the CHD cohort affecting proteins inside the GT-PPI (GT-PPI) or proteins outside the interactome (Non-Interactome). The white dot represents the median, the black lines the interquartile range (thick) and 1.5x the interquartile range (thin). P-values were determined using a two-sided Mann-Whitney-Wilcoxon test with Bonferroni correction; the number of asterisks indicate significance level (***p-value<0.001).

Since several of the GT-interactors with CHD-associated DNVs have previously been implicated in human cardiac malformations (Figure S3A) (Bouman et al., 2017; Chen et al., 2020; Ji et al., 2020; Jin et al., 2017; Jones et al., 2012; Maitra et al., 2010; Parisot et al., 2010; Pierpont et al., 2018; Thienpont et al., 2010), we sought to determine whether the enrichment was predominately driven by genes previously known to be involved in cardiac development or CHD by removing them from the dataset and repeating the permutation-based analysis (Table S5). We still found a significant enrichment in proteins harboring protein-altering DNVs from CHD probands in both GATA4 and TBX5 interactomes (adjusted OR G-PPI: 5.17; T-PPI: 3.38) (Figure 2B and Table S4). These results show that the integration of cardiac-specific TF-PPIs with CHD whole-exome sequencing studies can be used to unveil novel candidate genes enriched for mutations in CHD.

Most protein-protein interaction analyses are conducted in cell types that are convenient but less biologically relevant, and they often employ ectopic expression systems that sacrifice issues of stoichiometry. Our analysis, by contrast, was conducted in human cardiac progenitor cells and relied on the endogenous expression of TBX5 and GATA4. To assess the importance of a biologically relevant context for protein-protein interaction analysis, we generated GT-PPIs in kidney cells (HEK293T) over-expressing human GATA4 and TBX5 and subjected them to the same permutation analysis with the CHD and control cohorts (Figure S3B–S3D and Table S6A and S6B). There was no significant enrichment in proteins harboring CHD-associated protein-altering DNVs (adjusted OR G-PPI: 1.55; T-PPI: 1.04) (Figure 2B and Table S4). Although the HEK293’s GT-PPI (221 interactors) did not significantly differ in size compared to the CP’s GT-PPI (252 interactors), among all interactors found in kidney cells with GT-ectopic expression, only 20 GATA4 and 8 TBX5-interactors were verified by affinity purification and mass spectrometry of the endogenous cTFs in cardiac progenitors (Figure S3B-D and Table S6A&B). This result highlights the importance of endogenous tissue-specific protein-protein interactions in elucidating the genetic underpinnings of human diseases.

Having demonstrated that GT-PPIs were enriched in protein-altering variants found in the CHD cohort, we aimed to assess the likelihood that the GT-PPI variants contribute to disease. Using combined annotation-dependent depletion (CADD) scores, we found that GT-PPI protein-altering variants found in the CHD cases were more likely to be pathogenic than the rest of protein-altering DNVs in CHD cases outside the GT interactome (Figure 2C). These findings further validate the use of the interactomes of disease-associated TFs generated in biologically-relevant cell-types as a framework for identifying CHD candidate genes.

### GATA4:TBX5-Interactors with Protein-Altering DNVs Unveil CHD Candidate Genes with Characteristic Features of Disease Gene

We next asked whether the candidate CHD genes identified in the GT-PPI exhibited features that could increase their likelihood of causing disease. Extreme intolerance to loss of function (LoF) variation and haploinsufficiency are common features of genes associated with developmental disorders (Fuller et al., 2019). Remarkably, most candidate CHD genes in the PPI were extremely intolerant to LoF variation (probability of being intolerant to LoF (pLI) > 0.9) and exhibited significantly higher pLI and haploinsufficiency scores than genes outside the interactome with equivalent variants (Figure 3A and Figure S4A). Another feature of disease genes is an increased tendency for their products to interact with one another when their mutations result in similar phenotypes (Goh et al., 2007). Based on iRefIndex database information (Razick et al., 2008), the proteins encoded by our candidate genes had a higher connectivity degree with other proteins found to be mutated in the CHD cohort, as well as with a curated list of proteins involved in mouse/human cardiac malformations (Jin et al., 2017) than proteins outside the interactome with analogous variants (Figure 3B,C).

**Figure 3:**
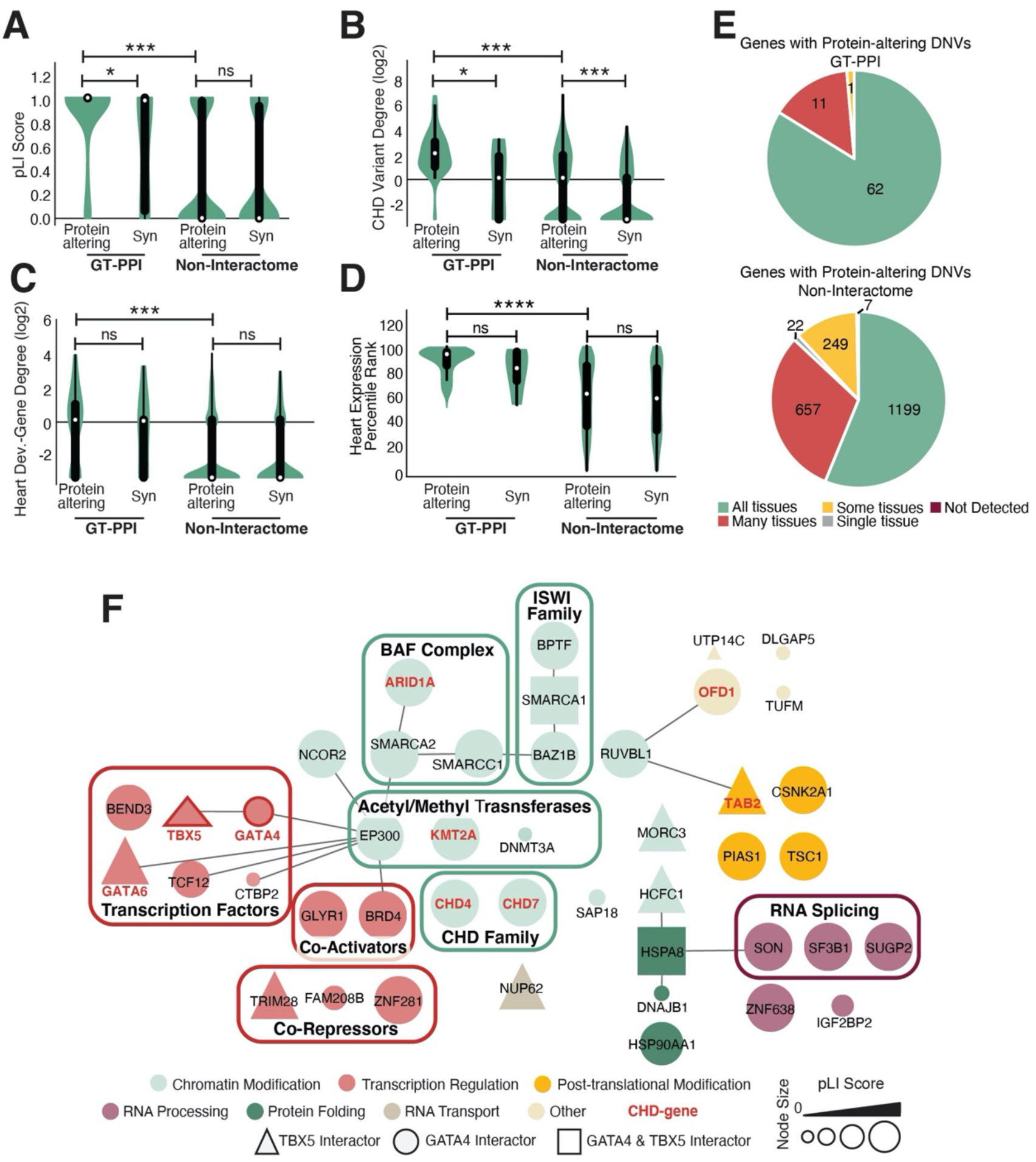
*De novo* variants in GATA4 and TBX5 interactomes exhibit features typical of disease genes. (A-D) Violin plots among GATA4 and TBX5 (GT) interactome proteins representing the distribution of (A) Intolerance to LoF (pLI Score); (B) degree of connectivity with all protein-altering DNVs found in the CHD cohort (CHD Variant Degree); (C) degree of connectivity with proteins encoded by a curated list of genes involved in mouse/human cardiac malformations (Heart Dev.-Gene Degree) (Jin et al., 2017); (D) expression percentile rank in the developing heart (E14.5) for Synonymous (Syn) or Protein-altering DNVs found in the CHD cohort and affecting proteins inside the GT interactome (GT-PPI) or outside the interactome (Non-Interactome). The white dot represents the median, the black lines the interquartile range (thick) and 1.5x the interquartile range (thin). P-values were determined using a two-sided Mann-Whitney-Wilcoxon test with Bonferroni correction; the number of asterisks indicate significance level (***p-value<0.001, *p-value<0.05). (E) Tissue expression distribution of interactome CHD candidate genes (GT-PPI) and non-interactome genes harboring CHD-associated protein-altering DNVs across the six Human Protein Atlas categories based on transcript detection (NX≥1) in all 37 analyzed tissues (See Methods). Detected in single: detected in a single tissue; Detected in some: detected in more than one but less than one third of tissues; Detected in many: detected in at least a third but not all tissues; Detected in all: detected in all tissues; Not detected. (F) Representation of interactome CHD candidate genes as a network after integration with PPI information from iRefIndex database. Nodes are colored based on manually annotated biological processes and specific protein families/complexes grouped in boxed areas. Node sizes reflect probability of Loss-of-function Intolerance (pLI) scores. Node shapes reflect belonging to TBX5 (triangle), GATA4 (circle) or GATA4&TBX5 (square) networks. Red highlights proteins encoded by genes involved in human CHD (Bouman et al., 2017; Chen et al., 2020; Ji et al., 2020; Jin et al., 2017; Jones et al., 2012; Maitra et al., 2010; Parisot et al., 2010; Pierpont et al., 2018; Thienpont et al., 2010). Edges represent protein-protein interactions from iRefIndex database (Razick et al., 2008).

GT-interactors with CHD protein-altering DNVs exhibited higher expression in the developing heart than genes with equivalent variants outside the GT-PPIs (Figure 3D). In addition, the CHD candidate genes we identified were generally expressed across most cell types in the developing heart and tissues in the human body (Figure 3E and Figure S4B–D) and largely affected functional modules relevant to chromatin biology (Figure 3F). Other biological processes with unexplored roles in CHD were affected, such as RNA splicing and protein folding (Figure 3F). Furthermore, many DNVs in GT-interactors were detected in probands suffering from CHD with extracardiac abnormalities and/or neurodevelopmental defects, but a sizeable number were also found in “isolated” CHD cases (Figure S4E and Table S7). This is consistent with previous studies from the Pediatric Cardiac Genomic Consortium that highlighted the functional relevance of high heart-expressed genes involved in chromatin modification and transcriptional regulation among cardiac malformations, with particular enrichment in CHD with coexisting neurodevelopmental defects (Homsy et al., 2015; Zaidi et al., 2013). Overall, our findings support the possibility that the candidate genes identified in the GT interactome are ubiquitously expressed genes with high heart-expression that are enriched for mutations that can cause different forms of CHD.

We next investigated the specific types of protein-altering *de novo* CHD variants corresponding to proteins in the GT-PPIs. Among the 252 proteins in the GT-PPI, 20 were encoded by genes harboring loss-of-function *de novo* variants present in CHD cases and 48 harboring missense CHD DNVs. The permutation-based statistical test detected strong enrichment within the GT-PPIs for both *de novo* loss-of-function (adjusted OR: 4.87) and missense DNVs (adjusted OR: 3.56) in CHD probands compared to equivalent genomic variation from the control cohort (Table S8). Loss-of-function *de novo* variants preferentially affected genes involved in human CHD (Table S7-8) (Garg et al., 2003; Ji et al., 2020; Jin et al., 2017; Jones et al., 2012; Maitra et al., 2010; Parisot et al., 2010), whereas the bulk of GT-PPI genes with CHD-missense DNVs had not previously been linked to cardiac development or CHD (Table S7-8). The contribution of *de novo* splice variants could not be determined due to their low counts in interactome genes from cases and controls (Table S8).

Collectively, these findings demonstrate that protein-altering *de novo* variants in GT-interactors found in CHD cases preferentially impact broadly expressed genes with high expression in the developing heart, and may therefore contribute to CHD outcomes through dosage effects. Of specific interest were GT-interactors harboring missense variants, which revealed candidate genes not previously implicated in congenital cardiac malformations.

### An Integrative Method for Scoring Variants Identifies Specific GT-Interactors as Strong Candidate Genes for CHD

The interpretation of missense variants remains an enormous challenge and requires methods to prioritize those that could substantially impact human phenotypes. The enrichment in missense DNVs for proteins yet unrelated to CHD in our interaction network highlighted the need for a strategy to interpret the likelihood that a rare variant in a given GT-PPI gene is contributing to disease. Previous studies from the Pediatric Cardiac Genomic Consortium used the gene’s ranked percentile of expression in the developing heart for filtering rare variants found in CHD cases (Jin et al., 2017; Zaidi et al., 2013). However, since most of the interactome CHD candidate genes were highly expressed in the developing heart (Figure 3D and Table S9), additional features were required to rank their likelihood of being pathogenic. Thus, we developed an integrative pipeline to calculate a potential pathogenesis score for the 48 missense DNVs mapped to our GT-PPI (Figure 4A). This score consolidates annotations from a combination of gene, variant and proband features relevant to protein-coding variants. Concretely, the scoring system prioritizes variants in GT-interactors based on gene or variant features found to be distinctive of the GT-PPI protein-altering DNVs, *e.g*., CADD score and pLI score (Figure 2D, Figure 3A–C and Figure S4D and S4F). At the proband level, it prioritizes variants detected in CHD cases without other reported DNVs or inherited variants in known CHD-genes within the same proband. The individual features were combined by rank sum and weighted where applicable (see Methods) (Figure 4A and Table S10). The scoring method, represented with respect to the genes’ percentile of expression of the developing heart, was able to separate published mutations causative of monogenic CHD (Basson et al., 1999; Furtado et al., 2017; Garg et al., 2003) from the few mutations known to cause oligogenic disease (Gifford et al., 2019) even when affecting the same gene; these are hereafter referred to as reference variants (Figure 4B). Furthermore, amidst the top-scored interactome variants, there were several proteins known to cause cardiac malformations, supporting the potential value of this prioritizing method (Figure 4B and Table S10).

**Figure 4:**
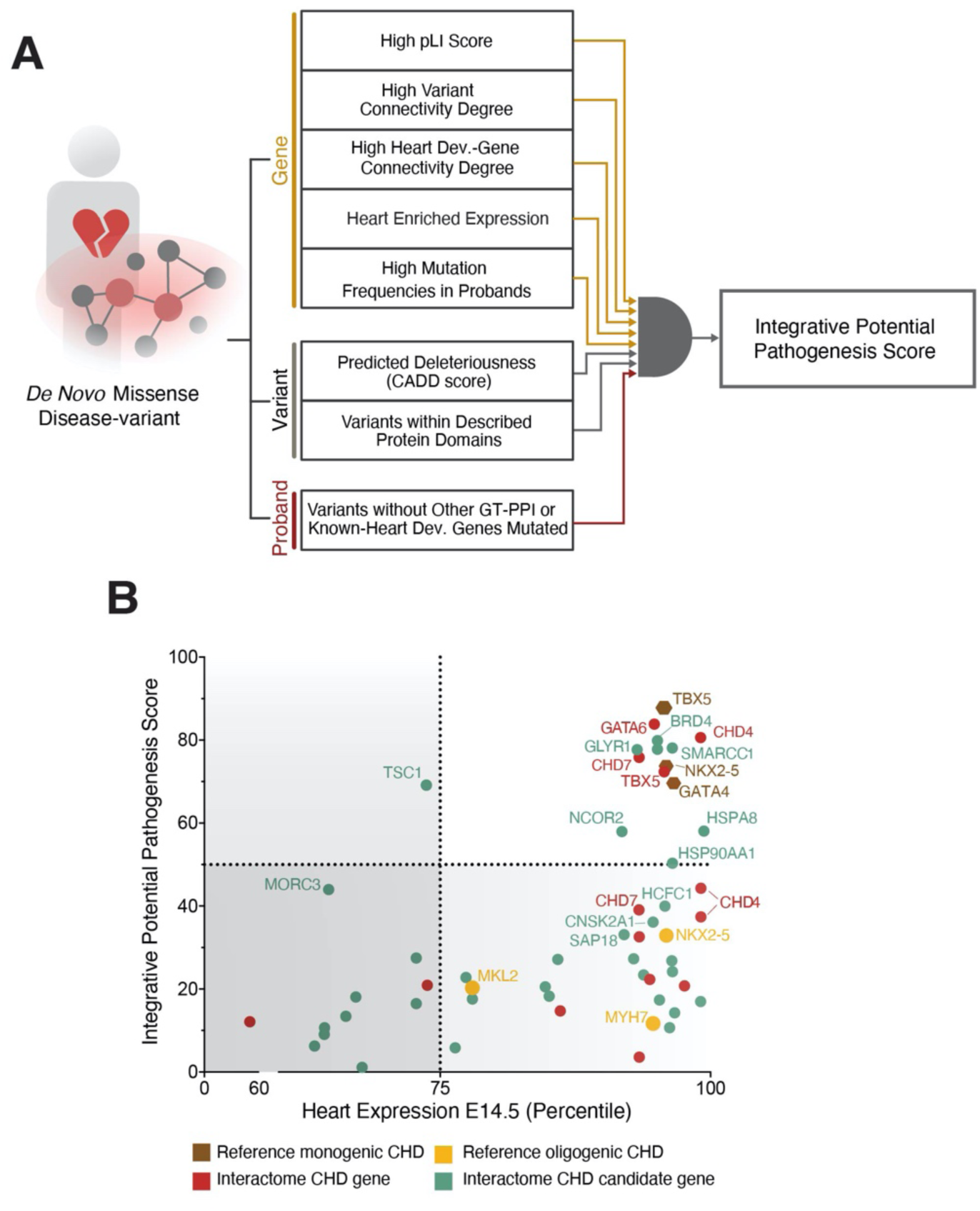
Integrative variant scoring devised to prioritize potential CHD-associated variants. (A) Integrative pathogenesis score developed from a combination of gene, variant and proband features distinctive of GT-PPI protein-altering variants. The indicated annotations were consolidated into a unique score by rank sum and weighted where applicable (see Methods). (B) Integrative pathogenesis scores for interactome missense DNVs in CHD genes (red) or in CHD unreported genes (green) plotted against the corresponding genes’ expression percentile rank in the developing heart (E14.5). Published mutations causing monogenic (grey) or oligogenic (orange) CHD are included as references.

The majority of the missense DNVs affected interactome proteins highly expressed in the developing heart, with only 25% occurring in GT-interactors outside the top quartile of expression; the lower expressed genes generally also exhibited low integrative potential pathogenesis scores, except for the tuberous sclerosis gene, *TSC1,* associated with cardiac rhabdomyomas (Hinton et al., 2014) (Figure 4B). On the other hand, missense DNVs in GT-interactors highly expressed in the developing heart clustered in two defined groups: a potentially highly pathogenic cluster of variants ranking close to the monogenic CHD references, and a group of variants close to the oligogenic CHD references. Among the missense DNVs in the first group, which we hypothesized to be more significant contributors, there were four variants in GT-interactors with previously described monogenic contribution to cardiac defects (*TBX5, GATA6, CHD4* and *CHD7*) and seven variants within proteins with yet undescribed functions in congenital heart malformations (*BRD4 x2, SMARCC1, GLYR1, HSPA8, NCOR2* and *HSP90AA1*) (Figure 4B and Table S10).

Three chromatin modifiers, BRD4, GLYR1 and SMARCC1, ranked the highest amongst the interactome proteins with unknown roles in CHD, in concordance with the observed increased burden of CHD-associated DNVs in genes involved in this process (Zaidi et al., 2013). These CHD candidate genes were detected as GATA4 interactors, and we validated each by co-immunoprecipitation (Figure S5A–S5D). While GLYR1 and SMARCC1 were previously unknown, the BRD4-GATA4 protein module was recently reported by our group in the regulation of cardiac mitochondrial homeostasis (Padmanabhan et al., 2020). In support of a BRD4 functional relevance in CHD, deletion of BRD4 during embryonic development (*Tnnt2-Cre; Brd4^flox/flox^)* resulted in embryonic lethality with signs of cardiac dysfunction (Padmanabhan et al., 2020). Although the specific contribution of SMARCC1 to CHD is yet uncertain, its encoded protein BAF155 is a component of the BAF complex, which orchestrates many aspects of heart development (Hota and Bruneau, 2016). The *GLYR1* DNV occurred in a patient with atrioventricular septal defects, left ventricle outflow tract obstruction and pulmonary stenosis, a spectrum of cardiac malformations observed in humans with *GATA4* mutations. However, the role of GLYR1 in most tissues, including the heart, remains unexplored.

### The CHD-Variant in *GLYR1* Impacts the Protein’s Structural Dynamics and Destabilizes its Physical Interaction with GATA4

GLYR1, also known as NDF, NPAC or NP60, is a chromatin reader involved in chromatin modification and regulation of gene expression through nucleosome demethylation (Fang et al., 2013; Fei et al., 2018; Fu et al., 2006; Marabelli et al., 2019; Yu et al., 2020). The *GLYR1* missense CHD *de novo* variant involved the substitution of a highly conserved proline at amino acid (aa) 496 for a leucine within the β-hydroxyacid dehydrogenase (β-HAD) domain, described to mediate the PPI between GLYR1 monomers (Marabelli et al., 2019; Montefiori et al., 2019). Since proline^496^ is located within a rigid loop enriched in aromatic residues connecting two tetramerization domains (Figure 5A–5C), we hypothesized the substitution of this rigid proline^496^ for a leucine would impact the structural dynamics of the GLYR1 β-HAD domain and therefore its ability to acquire certain functional states.

**Figure 5:**
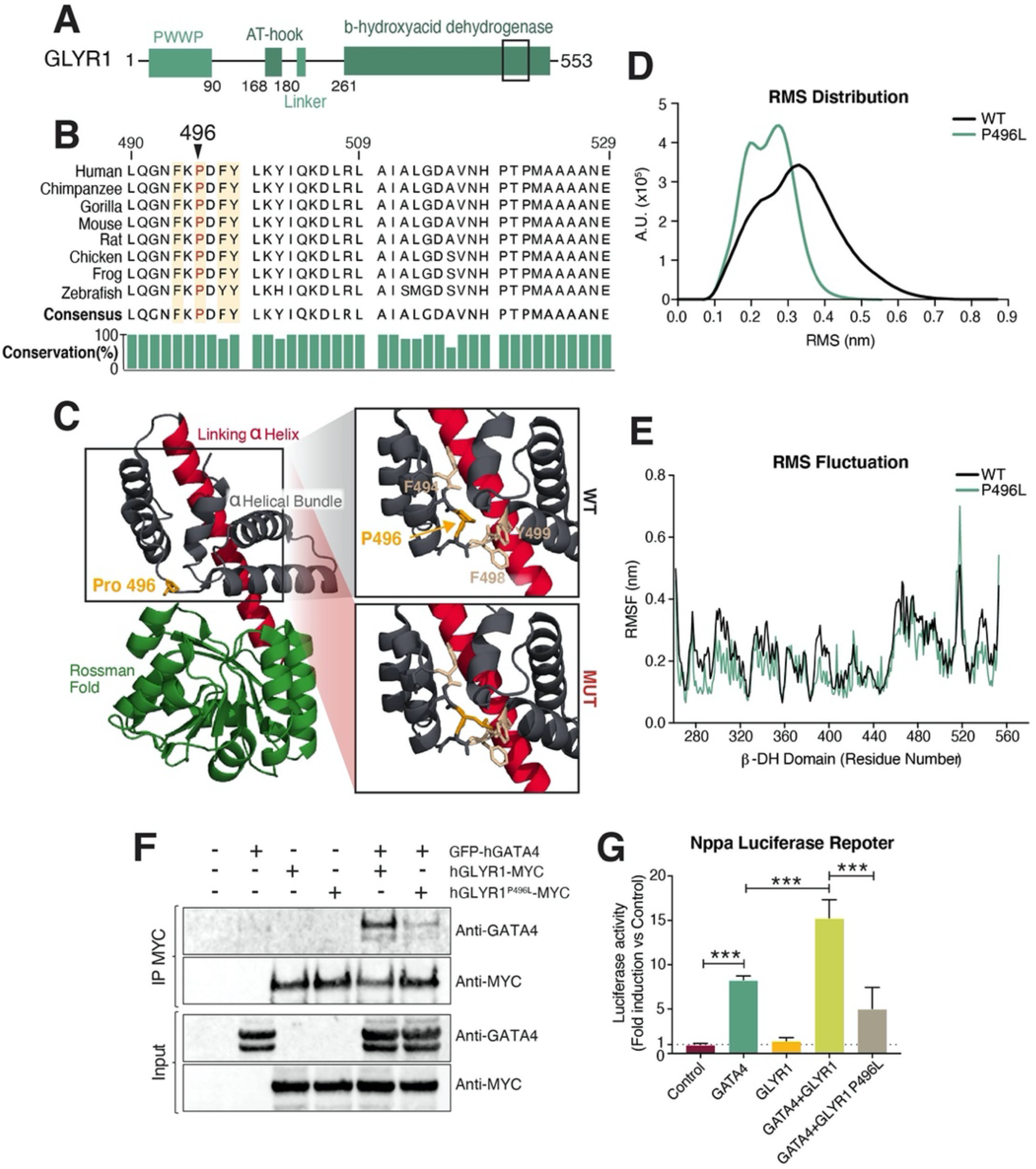
Functional impact of a highly scored CHD variant in GLYR1. (A) Simplified protein schematic depicting the domain organization of human GLYR1. Black rectangle indicates zoomed-in protein region in Figure 5B. (B) Protein sequence conservation across vertebrate species for the GLYR1 rigid loop region containing the CHD-associated P496L DNV. Amino acids 490-529, partially spanning exons 14 and 15 (490-495, 496-5229 respectively) in the *H. sapiens* sequence. (C) Subdivision of the GLYR1 dehydrogenase domain into the Rossman-fold globular domain (green), the linking α9-helix (red), and the α-helical bundle (dark blue). The mutated Proline^496^ is highlighted in orange. The right panels zoom into the WT and mutant forms of the rigid loop enriched in aromatic residues (beige) that contains the Proline 496 (orange). (D) Distribution of the root mean square deviation (RMSD) of frames visited during the trajectories from the reference state represented by the starting structure of the WT (black) and the P496L mutant (green) GLYR1 dehydrogenase domains within the measured time. (E) Residue flexibility analysis based on the standard deviations of the atomic positions in the simulations (RMSF) after fitting to the starting structure of the WT form (black) and the mutant (green) GLYR1 dehydrogenase domains. An overall lower flexibility (lower RMSF) of the mutant compared to the WT is most evident in the Rossman-fold domain (residues 262-437). (F) The ability of GLYR1 WT and P496L mutant to interact with GATA4 assessed by ectopic expression in HEK293 cells and immunoprecipitation (IP) of GLYR1-MYC followed by immunoblotting with the indicated antibodies. Enriched nuclear lysates prior to IP (Inputs) were set aside and analyzed in parallel with IP samples to verify similar protein ectopic expression levels across samples. (G) Luciferase reporter assay in HeLa cells showing activation of the luciferase reporter upon addition of plasmids encoding indicated proteins. (n=3 independent experiments). One-way ANOVA coupled with Tukey post hoc test: *** p-value <0.001.

Molecular dynamics (MD) computational simulations of the wild-type (GLYR1^WT^) and the CHD mutant (GLYR1^P496L^) β-HAD domains predicted the mutant β-HAD to explore a narrower set of structural conformations than the WT, as shown by the time-dependent evolution of the root mean square deviation (RMSD) of frames visited during the trajectories from the reference structure (Figure S6A). This result was confirmed by the distribution of the RMSD calculated for every pair of states sampled during the simulations (Figure 5D). Furthermore, GLYR1 structural dynamics at the local level, measured by the standard deviations of the atomic positions in the simulations (RMSF), indicated an overall lower flexibility of the P496L mutant compared to the WT, which was more evident in the Rossman-fold domain (262-437 aa) (Figure 5E). These data indicated that the P496L variant in GLYR1 induces significant differences in the structural dynamics of the β-HAD domain, at the global and local levels, predicting a general increase in the structural rigidity of this region in GLYR1.

The predicted increase in rigidity within the β-HAD domain could affect GLYR1’s capacity to adapt to interacting partner proteins through conformational selection. Co-immunoprecipitation assays demonstrated that the *GLYR1* P496L DNV destabilized its physical interaction with GATA4 (Figure 5F and Figure S6B). Since previous studies involved GLYR1 in transcriptional regulation (Fei et al., 2018; Yu et al., 2020), we probed whether GLYR1 co-regulates gene expression together with GATA4 by testing the ability of GLYR1 and GATA4 to transactivate the *Nppa*-luciferase reporter construct in transient transfection assays, a well-established assay of GATA4’s transcriptional activity (Garg et al., 2003; Hu et al., 2011; Knowlton et al., 1991). Transfection of GATA4 alone resulted in an approximate 8-fold activation of this reporter, whereas GLYR1 alone failed to induce luciferase activity. Co-transfection of GATA4 and GLYR1 increased reporter activity by approximately 15-fold, consistent with functional co-regulation. However, the synergistic activation induced by GLYR1 WT was attenuated by the P496L mutation (Figure 5G). Synergistic transactivation of the *Ccnd2*-luciferase reporter by GLYR1 and GATA4 was similarly reduced by the P496L mutation (Figure S6C). Overall, these findings demonstrate a protein-damaging effect of the GLYR1 missense *de novo* variant associated with CHD, which destabilized GLYR1’s physical interaction with GATA4 and impacted their transcriptional coregulation activity.

### GATA4 & GLYR1 Co-bind a Defined Set of Heart Development Genes and Co-Regulate their Expression

GLYR1 localizes within chromatin regions rich in histone H3 trimethylated on Lys36 (H3K36me3) at actively transcribed gene bodies to regulate transcription elongation (Fang et al., 2013; Fei et al., 2018; Marabelli et al., 2019; Yu et al., 2020). However, knowledge about how GLYR1 is recruited to specific loci or its function in homeostasis and disease is limited. We analyzed its genome-wide occupancy during cardiomyocyte differentiation together with H3K36me3 genome-wide distribution and gene expression activity. ChIP-sequencing (ChIPseq) in hiPSCs and CPs revealed dynamic relocalization of GLYR1 from hiPSCs to CPs. Although genomic regions bound by GLYR1 in hiPSCs were largely maintained in CPs, we detected ∼3500 differentially bound genes (FDR<0.1) between the two stages (Figure 6A and Figure S7A). K-means clustering of genes differentially bound by GLYR1 based on the three measured variables—GLYR1 ChIPseq, H3K36me3 ChIPseq, and RNA expression—highlighted GLYR1 recruitment to ∼3000 gene bodies upon differentiation of hiPSCs to CPs (Clusters 1 and 2). Gene ontology (GO) analysis revealed that gene programs associated with heart development were enriched in Cluster 2, which showed the highest levels of GLYR1 ChIP signal in CPs, whereas Cluster 1 was enriched for genes involved in general cellular processes. On the other hand, Cluster 3 contained ∼500 GLYR1-bound genes in hiPSCs and lost in CPs, mainly associated with cell cycle and ribosome biogenesis terms (Figure 6A and Table S11 and S12). Overall, GLYR1 preferentially bound to transcribed regions of active genes and co-localized with H3K36me3 (Figure 6A and Figure S7A–D). Interestingly, GLYR1 only occupied a fraction of the genes up-regulated in CPs and marked with H3K36me3 (Figure S7A–S7C), suggesting that GLYR1 binds a very discrete set of loci marked by H3K36me3. These results revealed dynamic recruitment of GLYR1 to a discrete set of cardiac genes during cardiomyocyte differentiation.

**Figure 6:**
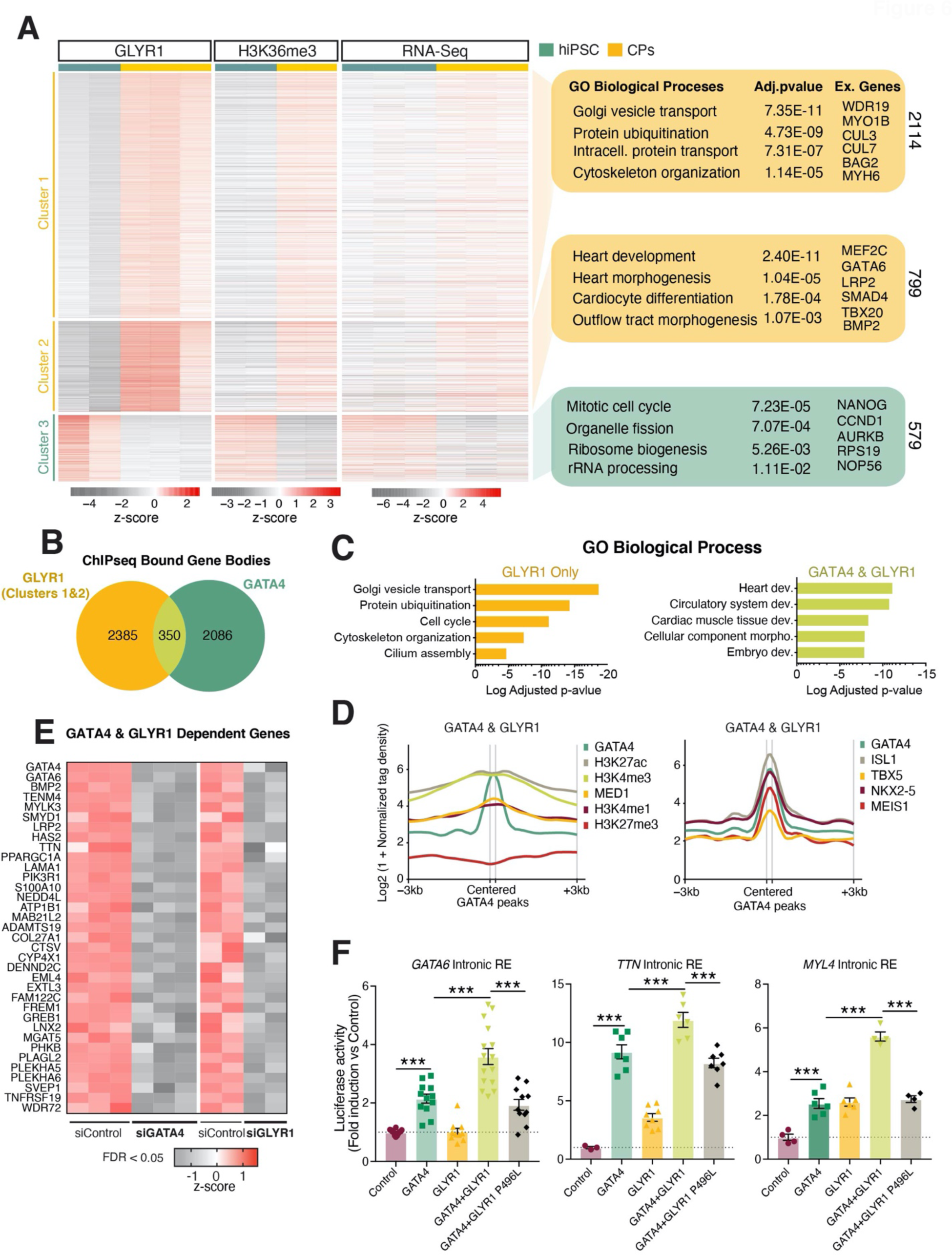
GATA4-associated and-independent roles for GLYR1 in transcription regulation during cardiomyocyte differentiation. (A) Genes differentially bound by GLYR1 (FDR<0.1) between hiPSCs and cardiac progenitors (CPs) subjected to k-means clustering based on the three indicated variables in hiPSC and CP stages: GLYR1 ChIPseq signal, H3K36me3 ChIPseq signal and gene expression levels. Statistically enriched GO Biological Process terms and example genes for each cluster on the right panel. hiPSC GLYR1, hiPSC and CP H3K36me3 ChIPseq (n=2); CPs GLYR1 (n=3), publicly available RNAseq hiPSC and CP (GSE137920; n=3). (B) Venn Diagram showing the overlap of genes bound by GLYR1 in CPs from Clusters 1 & 2 (FDR<0.1, LogFC>0.5) with genes occupied by GATA4 inside the gene body window (1^st^ intron-TES) where GLYR1 can be found. The significance of the overlap estimated using the Fisher.Exact function is p-value = 2.865e^-16.^ The odds of Gata4 binding to bodies of genes with enriched Glyr1 signal is 1.69 times the odds of no Gata4 binding in bodies of genes enriched with Glyr1 signal. (C) Gene Ontology enrichment analysis of biological process for GATA4 & GLYR1-bound genes or GLYR1-only bound genes. Dev, development. (D) GATA4 & GLYR1 co-bound genes that were significantly downregulated (FDR<0.05, LogFC<-0.25) upon knockdown of either GATA4 or GLYR1 at the cardiac progenitor stage. Cells were transfected with the corresponding siRNAs at day 4 of differentiation and cardiac progenitors collected 72h later for RNAseq. (E) Metagene plots for GATA4 & GLYR1 co-bound genes plotting the normalized ChIPseq signal for the indicated histone marks (public available data GSE85631 and GSM2047027) and other cardiac transcription factors centered on GATA4 peaks within the gene body (1^st^ Intron-TES). One representative replicate plotted, all replicates in Figure S6J. (F) Transcriptional activity of three putative intronic regulatory elements co-bound by GATA4 & GLYR1 in the presence of indicated regulatory proteins. Intronic sequences were cloned into Luciferase reporters and luciferase activity was assayed in HeLa cells upon addition of plasmids encoding the indicated proteins. Equal amount of total transfected DNA per condition was adjusted with empty vector. (n= 3 independent experiments). One-way ANOVA coupled with Tukey post hoc test: *** p-value <0.001.

In CPs, as described in other cell types (Fei et al., 2018; Yu et al., 2020), GLYR1 broadly occupied gene bodies (Figure S7D), from the first intron to the transcription end site (TES) on average. On the other hand, GATA4 preferentially occupied distal regulatory elements, though some peaks are also found at introns inside gene bodies, similar to GLYR1 (Figure S7E). To investigate GATA4-GLYR1 genomic co-occupancy in CPs, we overlapped the genes where GLYR1 was recruited in CPs (GLYR1^CP^: clusters 1-2, FDR<0.1 and Log2FC>0.5) with genes bound by GATA4 within the gene body window where GLYR1 typically binds (1^st^ Intron-TES). This analysis found a statistically significant overlap between GLYR1^CP^ and GATA4-bound gene bodies (Fisher exact p-value = 2.865E^-16^; OR: 1.69), identifying a defined subset of GATA4 and GLYR1-bound genes enriched in heart development GO terms (Figure 6B and 6C and Table S12 and S13). On the other hand, GLYR1-occupied sites that were not bound by GATA4 mapped to genes controlling general processes such as vesicle transport and protein ubiquitination. Among the gene bodies occupied by GATA4 only, approximately half were down-regulated from hiPSCs to CPs and enriched in neurogenesis-related terms, while the other half were up-regulated and associated with circulatory system development and cell adhesion (Figure 6C, Figure S7F and Table S13–15).

To directly evaluate if GATA4 and GLYR1 regulate the transcript levels of the genes they co-occupy, we analyzed the effect of silencing GATA4 or GLYR1 on the expression of these genes in CPs by bulk RNAseq (Table S16 and S17). GLYR1 silencing led to reduced expression of more than 800 genes associated with embryonic development and heart development terms compared to a control siRNA, which suggested a functional relevance for GLYR1 in the transcriptional regulation of the cardiomyocyte differentiation process (Figure S7G and Table S17 and S18). At loci co-bound by GATA4 and GLYR1, gene expression was 10 times more likely to be significantly down-regulated by either GATA4 or GLYR1 knockdown compared to those not co-bound (Figure S7H). Several co-occupied and co-regulated loci (LRP2, HAS2, TEMN4, SMYD1, MAB21L2, TTN, GATA4, GATA6) are involved in human or mouse cardiac malformations (Baardman et al., 2016; Camenisch et al., 2000; Nakamura et al., 2013; Park et al., 2010; Rasmussen et al., 2015; Saito et al., 2012; Theis et al., 2019; Zhu et al., 2014) (Figure 6D). The observation that GATA4 mainly occupies intronic regions within GATA4:GLYR1-bound gene bodies led us to ask whether GATA4 was binding intronic regulatory elements at co-occupied genes. To answer this, we examined features characteristic of active or repressed gene regulatory elements (Akerberg et al., 2019; Kimura, 2013) among the GATA4-occupied sites. GATA4 occupancy within GATA4:GLYR1-bound gene bodies co-localized with high levels of marks associated with active regulatory elements (H3K27ac, H3K4me3, H3K4me1, MED1) as well as with other cardiac TFs, but with undetectable levels of the repressive mark H3K27me3 (Figure 6E and Figure S7I). These findings raised the possibility that GATA4 and GLYR1 positively regulate cardiac gene expression by co-binding and activating intronic regulatory elements.

To investigate this hypothesis, we cloned several intronic regions with the features described above into a luciferase reporter vector under control of a minimal promoter and tested the ability of GLYR1 and GATA4 to transactivate the reporter. Transfection of GATA4 alone resulted in activation of these reporters, whereas GLYR1 alone induced luciferase activity of two out of three tested reporters, indicating that these intronic locations could function as REs (Figure 6F). Importantly, co-transfection of GATA4 and GLYR1 increased reporter activity, consistent with functional co-regulation. Remarkably, the synergistic/additive activation induced by GLYR1 WT was strongly attenuated in the context of the GLYR1 P496L mutation (Figure 6F). These findings suggest GATA4 recruits GLYR1 to co-regulate the expression of a discrete set of genes essential for heart development and disruption of this interaction by the *GLYR1* P496L missense DNV associated with CHD affects this co-regulation.

## DISCUSSION

Here, we integrated an analysis of the protein-protein interaction network of CHD-associated TFs with human whole-exome sequencing data to inform the genetic underpinnings of CHD. An unbiased PPI reconstruction for two essential TFs in cardiac development that cause CHD, GATA4 and TBX5, identified known and previously unreported functional relationships for these well-studied TFs. *De novo* mutations in GT-PPIs occurred with significantly greater frequency in CHD patients than healthy controls. Additionally, a consolidative computational framework devised to prioritize variants in GT-interacting proteins identified numerous candidate disease genes, including GLYR1, a ubiquitously expressed epigenetic reader. GLYR1 widely co-occupied cardiac regulatory elements with GATA4, and the *GLYR1* disease variant disrupted the GATA4 interaction and co-activation of cardiac developmental genes. These findings indicate that the use of tissue-and disease-specific PPIs may partially overcome the genetic heterogeneity of CHDs and help prioritize the potential impact of de novo missense variants present in disease.

### Integration of Tissue-specific TF-PPIs with Human Variant Data Highlights Disease Mechanisms

GT-PPIs were enriched in proteins harboring likely deleterious CHD-associated DNVs, validating the use of CHD-associated TFs-PPIs generated in disease-relevant cell-types as a framework for identifying CHD candidate genes. Unlike other approaches that rely on tissue-specific expression as a crucial criteria for narrowing potential genetic contributors to disease (Bryois et al., 2020), this strategy allows us to capture ubiquitously expressed CHD candidate genes that might have tissue-specific effects thanks to their interaction with tissue-enriched factors. This is of great importance as the majority of disease genes have been shown to be broadly expressed across multiple human tissues.

Previous studies have also applied gene-set enrichment strategies that calculate the mutation burden in defined groups of proteins, based on known disease genes or functional modules within variant-gene-networks reconstructed from consolidated PPI databases (Caldera et al., 2017; Izarzugaza et al., 2019; Magger et al., 2012; Zaidi et al., 2013). However, these studies had strong dependence on a curated list of already established disease candidate genes or on publicly available PPI information collected from various cell types, often employing ectopic expression systems. Our findings highlighted the importance of reconstructing PPIs in a disease-relevant context and with endogenous proteins as baits in order to maximize the ability to infer causal genes. Advances in proteomic technologies and the increased availability of tissue-specific public data will be essential to facilitate the broad and high-throughput deployment of network-based strategies. These strategies may not be limited to protein interaction networks and coding variants, as networks of regulatory elements based on ChIP-sequencing could also be integrated with data sets of non-coding variants.

Unlike single-gene enrichment approaches, the network-enrichment analysis allows the detection of rare CHD candidate genes, but it does so without resolving the relative contributions of specific variants. Hence, downstream prioritization of candidate disease genes is needed to rank likelihood of specific variants contributing to CHD. For this purpose, we developed an integrative scoring method that combines commonly used disease-variant prioritization metrics, including diverse and complementary biological information at the gene, variant and proband levels. The integration of proband genomic information regarding the co-occurrence of rare inherited and/or DNVs in known CHD genes, with metrics that predict variant deleteriousness and gene-level parameters that calculate their likelihood of being CHD risk genes, allowed separation of potential highly penetrant variants from possible modifier variants contributing to polygenic CHD. Although we focused our functional studies on variants clustering with known monogenic genes, these data present numerous avenues to study the underpinnings of oligogenic and monogenic types of congenital cardiac malformations presenting alone or in combination with extra-cardiac defects. Functional investigation will be needed to test whether the identified CHD candidate genes are essential in heart development and to determine the causal nature of the associated variants. In the future, high-throughput screening methods, such as the integrative PPI-genetic variant scoring pipeline, will aid in assessing the vast genomic variation catalogue provided by the increasing number of large-scale sequencing studies.

### GLYR1 Co-regulates Heart Development Genes with GATA4 and Is Mutated in CHD

The integrative proteomics and human genetics approach revealed GLYR1 as a previously unreported GATA4 interactor in CPs that constitutes a strong candidate gene for CHD. Leveraging computational simulations and biochemical assays, we demonstrated that the GLYR1 P496L variant impacted the protein structural dynamics and reduced its interaction with GATA4. This raised the possibility that destabilization of tissue-specific PPIs could result in cardiac-restricted phenotypic manifestation associated with the mutation of a ubiquitously expressed chromatin reader. As yet unstudied is the possibility that this missense DNV in GLYR1 may cause broader effects on the GLYR1 interactome network or its ability to adopt quaternary structures that could contribute to the disease phenotype. Despite its ubiquitous expression, the functional importance of GLYR1 protein across tissues in development and disease to date remains unexplored. To our knowledge, this work highlighted for the first time a role for GLYR1 in the transcriptional regulation of essential genes in heart development during cardiomyocyte differentiation.

Genome-wide GLYR1 occupancy interrogation in hiPSCs and CPs confirmed a cell-type specific dynamic binding of GLYR1 to gene bodies marked by H3K36me3, where GLYR1 recruitment correlated with active transcription as expected (Fei et al., 2018; Yu et al., 2020). The GLYR1 localization to only a fraction of the H3K36me3-enriched regions suggested specificity of its DNA occupancy. Our work indicates that during cardiomyocyte differentiation, GATA4 physical interaction with GLYR1 may be one of the mechanisms explaining how GLYR1 can bind a specific subset of heart development genes. However, only a fraction of the heart development genes to which GLYR1 is recruited upon differentiation of hiPSCs to cardiac progenitors is co-bound by GATA4, suggesting that GLYR1 might also interact with other cardiac-enriched factors.

Overall, this work has identified novel interactors of TFs essential for cardiac development, provided a ranked list of candidate disease genes with variants potentially contributing to CHD, and revealed novel biology of gene regulation related to cardiac disease. Notably, this tissue-and disease-specific TF network-based approach can be applied to other genetic disorders for which large-scale sequencing data is available to highlight disease mechanisms and provide a powerful filter for interrogating the genetic basis of disease.

## ACKNOWLEDGMENTS

We thank the Srivastava laboratory and Gladstone colleagues for critical discussions and feedback; Guadalupe Sabio and Mauro Costa for critical reading of the manuscript; B. Taylor for editorial assistance; Jim Kadonaga for kindly sharing GLYR1 anti-serum; the Gladstone Genomics Core, Bioinformatic Core, Stem Cell Core and Flow Cytometry Core for their technical expertise and the Gladstone Animal Facility for support with mouse colony maintenance; David E. Gordon for sharing an optimized CRISPR/Cas9 RNPs hiPSCs knockout generation protocol. We thank Francoise Chanut for manuscript editorial support and Ana Catarina Silva for helping with figure editing and design.

## AUTHOR CONTRIBUTIONS

B.G.T., K.S.P., and D.S. conceived and directed the study with input from B.G.B., B.R.C., B.L.B., and N.J.K. as part of an NIH/NHLBI-sponsored Program Project Grant and B.D.G. as part of the Pediatric Cardiac Genomics Consortium. D.R.B. and B.G.T generated the GATA4-KO hiPSC line and B. Cole and B.G.T. generated the TBX5-KO hiPSC line. D.R.B. and B.G.T. performed WT, GATA4 and TBX5 knockout CP and CM differentiations and characterization. B.C., R.H. and B.G.T. defined the appropriate affinity-purification strategy for cTFs. B.G.T. performed GATA4 and TBX5 (GT) affinity purification and sample preparation for mass spectrometry and GT-PPI classification. M.M. prepared the APMS buffers and performed the desalting and lyophilization of the APMS samples. M.P. and K.S.P. performed the APMS statistical analysis and M.P., K.S.P. and B.G.T. the PPI filtering. M.P. and K.S.P. performed GT-PPI variant enrichment and disease diagnosis association analysis. M.P. and B.G.T. performed interactome gene features analysis and developed the integrative pathogenicity scoring method. F.F. performed the GLYR1 protein sequence alignment, GLYR1 protein structure modeling and recombinant DNA cloning. E.M. and G.C. performed the molecular dynamics GLYR1 simulations. F.F. and B.G.T. performed GATA4 and GLYR1 silencing in CPs and luciferase reporter assays. K.S. generated the cTFs ChIPseq data. B.G.T. and M.A. performed GLYR1 and H3K36me3 ChIPseq. R.T., K.C. and C.G. performed RNAseq and ChIPseq computational analysis.

## SI FIGURE LEGENDS

**Figure S1.**
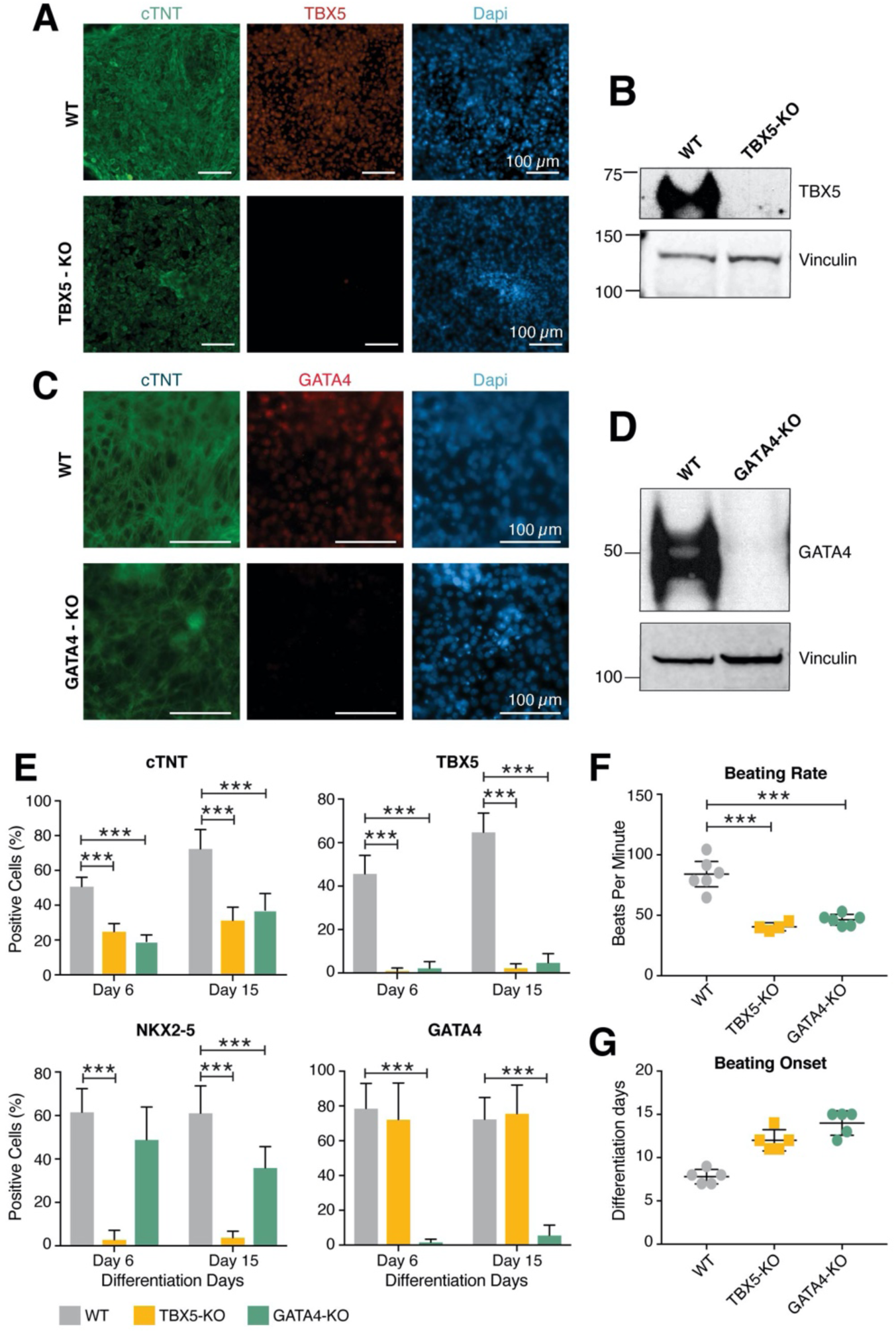
Differentiation of GATA4-KO and TBX5-KO hiPSC clonal lines into cardiomyocytes. Related to Figure 1. (A) Representative immunostaining micrographs for cTNT (green), TBX5 (red) or DAPI (blue) in WT or TBX5-KO hiPSC-derived cardiomyocytes (CM) at day 15 of differentiation. Scale (100μm). (B) Immunoprecipitation of TBX5 from enriched nuclear lysates of WT or TBX5-KO hiPSC-derived cardiac progenitors (CPs; differentiation day 6), followed by immunoblotting with anti-TBX5 or anti-vinculin antibodies. (C) Representative immunostaining micrographs for cTNT (green), GATA4 (red) or DAPI (blue) in WT or GATA4-KO hiPSC-derived cardiomyocytes (CM) at day 15 of differentiation. (D) Immunoprecipitation of GATA4 from enriched nuclear lysates of WT or GATA4-KO hiPSC-derived cardiac progenitors (CPs; differentiation day 6), followed by immunoblotting with anti-GATA4 or anti-vinculin antibodies (E) Percentage of cells positive for the indicated proteins at the CP (day 6) and CM (day 15) stages of differentiation as measured by flow cytometry. (n= 10-4) (F) Beating rates of the WT, TBX5-KO and GATA4-KO CMs as measured by Pulse automated measurement video analysis. (n=4-6) (G) Beating onset for WT, TBX5-KO and GATA4-KO CMs. (n=5) For E and F One-way ANOVA coupled with Tukey post hoc test: ***= p-value<0.001.

**Figure S2.**
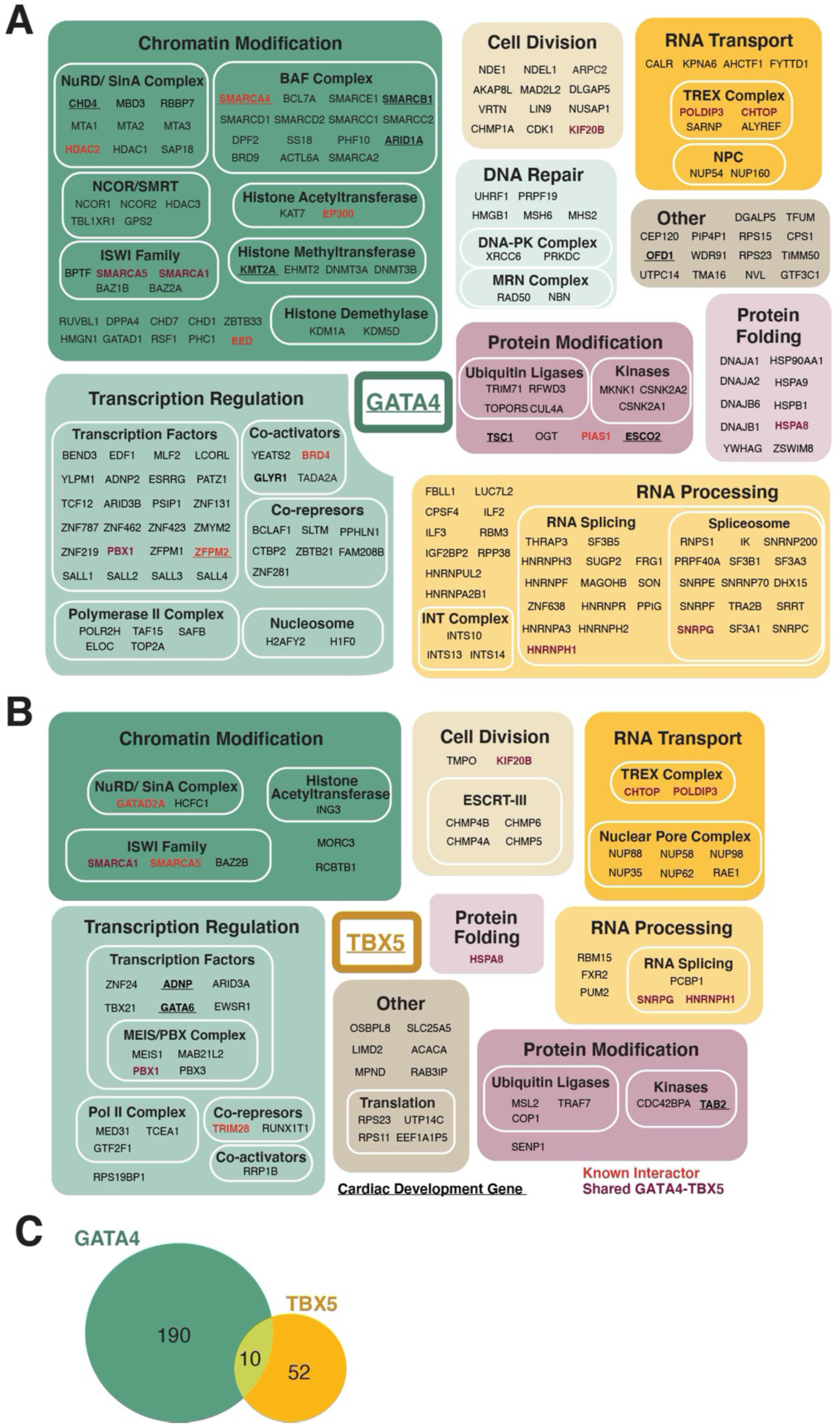
Complete GATA4 and TBX5 PPIs in hiPSC-derived cardiac progenitors. Related to Figure 1. (A) GATA4-PPI or (B) TBX5-PPI. Interactors were manually annotated for biological processes and protein complexes based on literature available. Boxed areas are roughly proportional to the number of interactors they represent. Enriched proteins with a Bayesian false discovery rate (BFDR)<0.001 for GATA4-PPI and BFDR<0.05 for TBX5-PPI are shown. Proteins interacting with both GATA4 and TBX5, previously reported interactors, and genes involved in mouse/human cardiac development (Jin et al., 2017) are highlighted in purple, red, and underline, respectively. 3-4 replicates from independent differentiations were analyzed per condition. (C) Venn diagram representing the overlap of GATA4 and TBX5 PPIs generated in CPs.

**Figure S3.**
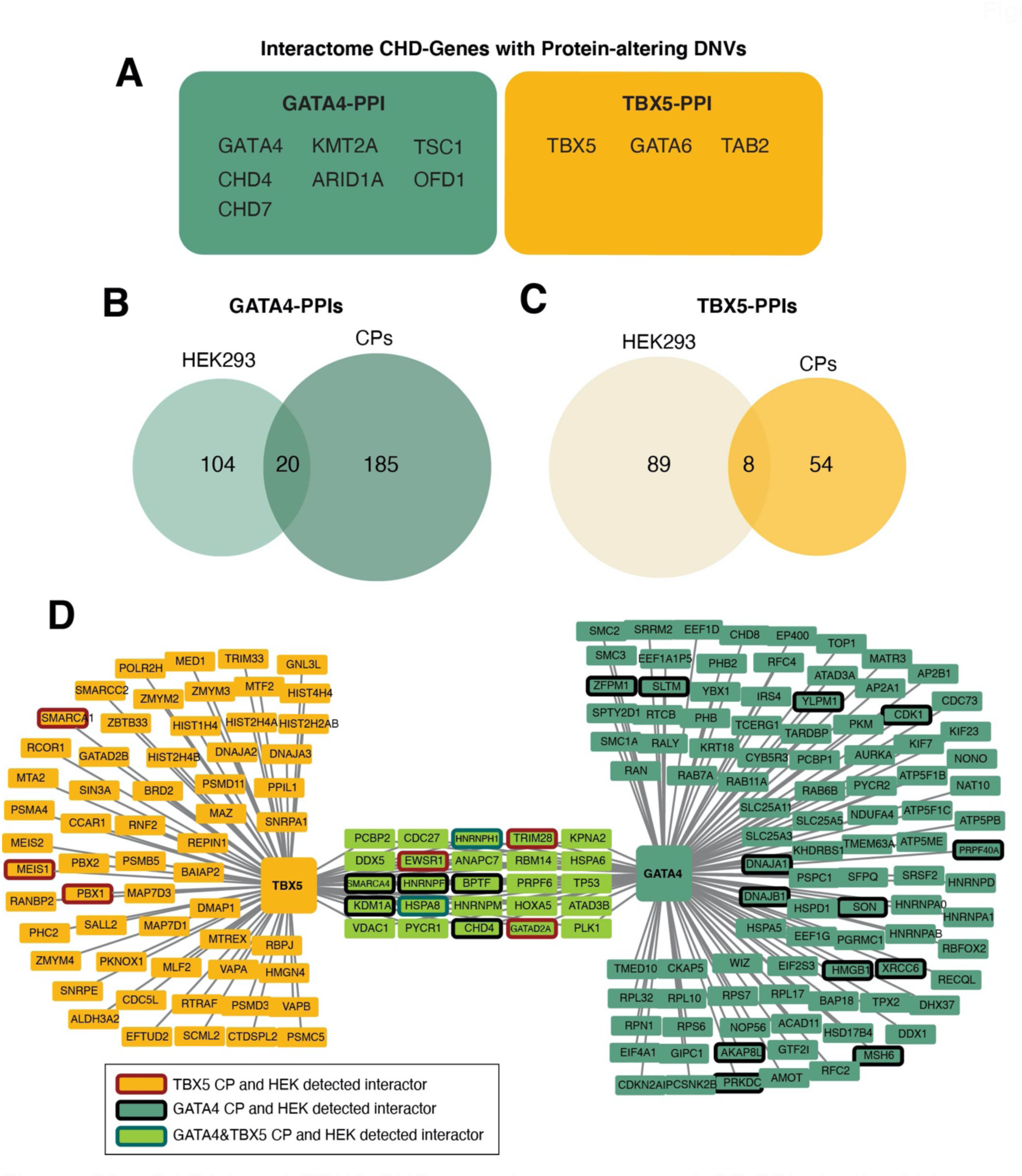
GATA4 and TBX5 CHD-gene interactors and GT-PPIs in the kidney cell line HEK293. Related to Figure 2. (A) GT-interactors with CHD-associated DNVs previously implicated in human cardiac malformations (Bouman et al., 2017; Chen et al., 2020; Jin et al., 2017; Jones et al., 2012; Maitra et al., 2010; Parisot et al., 2010; Pierpont et al., 2018; Thienpont et al., 2010). (B-C) Venn diagram representing the overlap of the GATA4 or TBX5 PPIs between hiPS cell-derived CPs and HEK293 cells. (D) GT-PPI reconstructed in HEK293 kidney cells. FLAG tagged GATA4 or TBX5 proteins were ectopically expressed in HEK293T cells and the cells collected 48h after transfection; an empty vector was used as negative control. Nuclear-enriched lysates treated with benzonase (DNase/RNase enzyme) were subjected to affinity purification (AP) with anti-FLAG antibodies. For each AP condition, replicates from three independent transfections were analyzed by mass spectrometry (LC/MS). AP-MS results from the negative controls were used to remove antibody-specific background from the experimental samples’ signal; data were subjected to the same filtering steps as the CP AP-MS data to identify high-confidence GATA4 and TBX5 PPIs. Enriched proteins with a BFDR<0.05 are represented in the network. CP and HEK293 overlapping TBX5, GATA4 and TBX5 & GATA4 interactors are highlighted with a colored node border in brown, black and green respectively.

**Figure S4.**
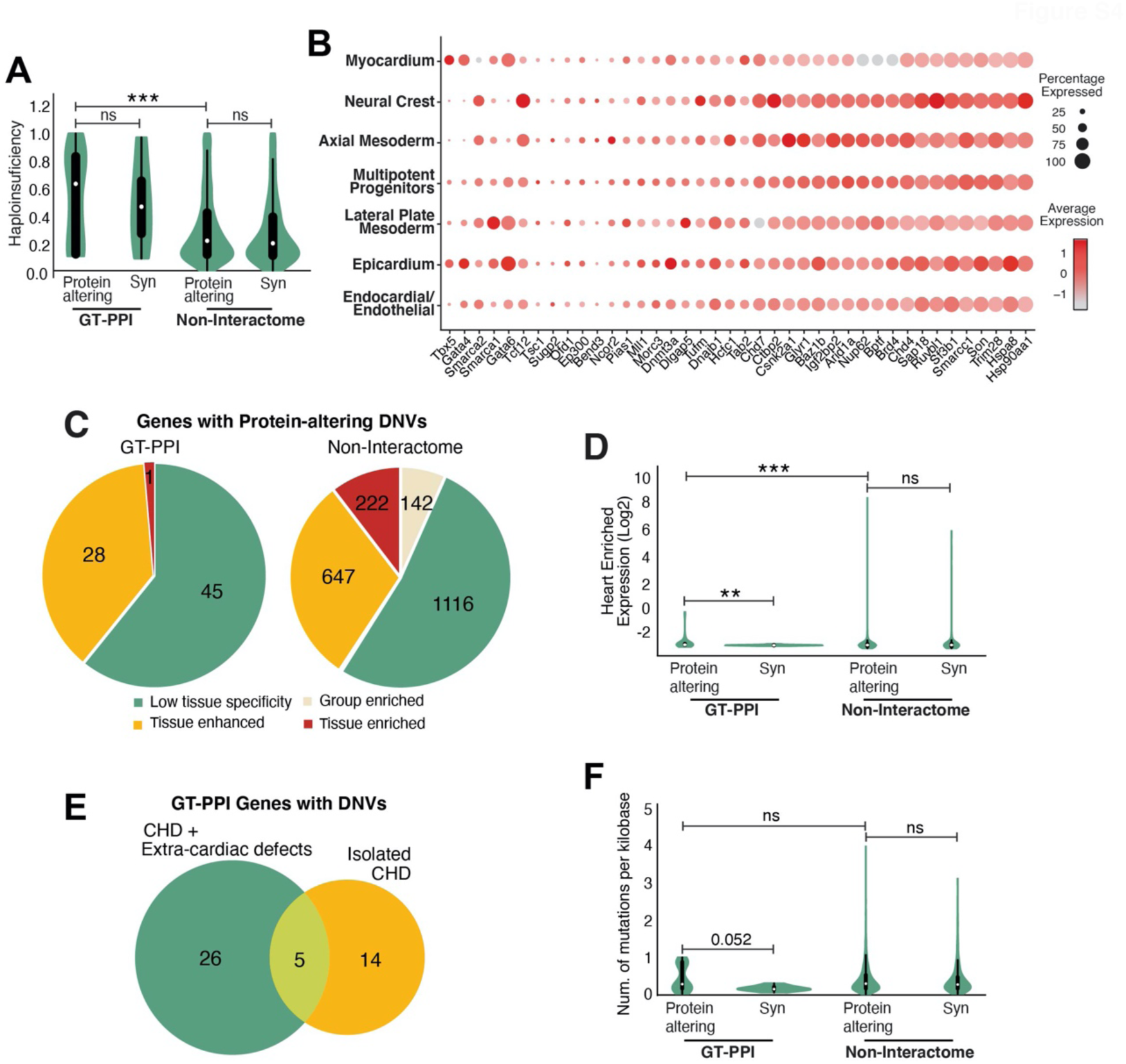
Features of CHD candidate genes in the GATA4-TBX5 interactome. Related to Figure 3. (A) Violin plot of the haploinsufficiency scores for synonymous (Syn) or protein-altering DNVs found in the CHD cohort and affecting proteins inside the GT interactome (GT-PPI) compared to outside the interactome (Non-Interactome). The white dot represents the median, the black lines the interquartile range (thick) and 1.5x the interquartile range (thin). P-values were determined using a two-sided Mann-Whitney-Wilcoxon test with Bonferroni correction; the number of asterisks indicate significance level (***p-value<0.001). (B) Dot plot representing the expression patterns of interactome genes harboring CHD-associated protein-altering DNVs in the mouse developing heart (average of E7.75, E8.25 and E9.25) based on published single-cell RNAseq data (de Soysa et al., 2019). The size of the dot indicates the percentage of cells expressing that gene within a cluster and the color indicates the average expression level of that gene within a cluster. (C) Distribution of GT-PPI and Non-Interactome genes harboring CHD-associated protein-altering DNVs across the five Human Protein Atlas categories based on transcript specificity in 37 analyzed tissues (See Methods). Tissue enriched: At least four-fold higher mRNA level in a particular tissue compared to any other tissues; Group enriched: At least four-fold higher average mRNA level in a group of 2-5 tissues compared to any other tissue; Tissue enhanced: At least four-fold higher mRNA level in a particular tissue compared to the average level in all other tissues; Low tissue specificity: detected and not within the other categories; Non detected. (D) Violin plot representing the distribution of Heart Enriched Expression (Log_2_ Heart GTEX RPKM/ Average RPKM in 18 non-heart tissues) for synonymous (Syn) and protein-altering DNVs found in the CHD cohort and affecting proteins inside the GT interactome (GT-PPI) or outside the interactome (Non-Interactome). The white dot represents the median, the black lines the interquartile range (thick) and 1.5x the interquartile range (thin). P-values were determined using a two-sided Mann-Whitney-Wilcoxon test with Bonferroni correction; the number of asterisks indicate significance level (**p-value<0.01, ***p-value<0.001). (E) Venn diagram representing the number of interactome genes with protein-altering DNVs found in probands suffering from “isolated CHD”, CHD with concomitant extra-cardiac defects (extracardiac abnormalities and/or neurodevelopmental defects), or in both types of CHD. (F) Number of mutations per kilobase, based on the number of mutations per gene reported by the *PCGC* (REF) corrected by the gene’s length, for synonymous (Syn) and protein-altering DNVs found in the CHD cohort and affecting proteins inside the GT interactome (GT-PPI) or outside the interactome (Non-Interactome). The white dot represents the median, the black lines the interquartile range (thick) and 1.5x the interquartile range (thin).

**Figure S5.**
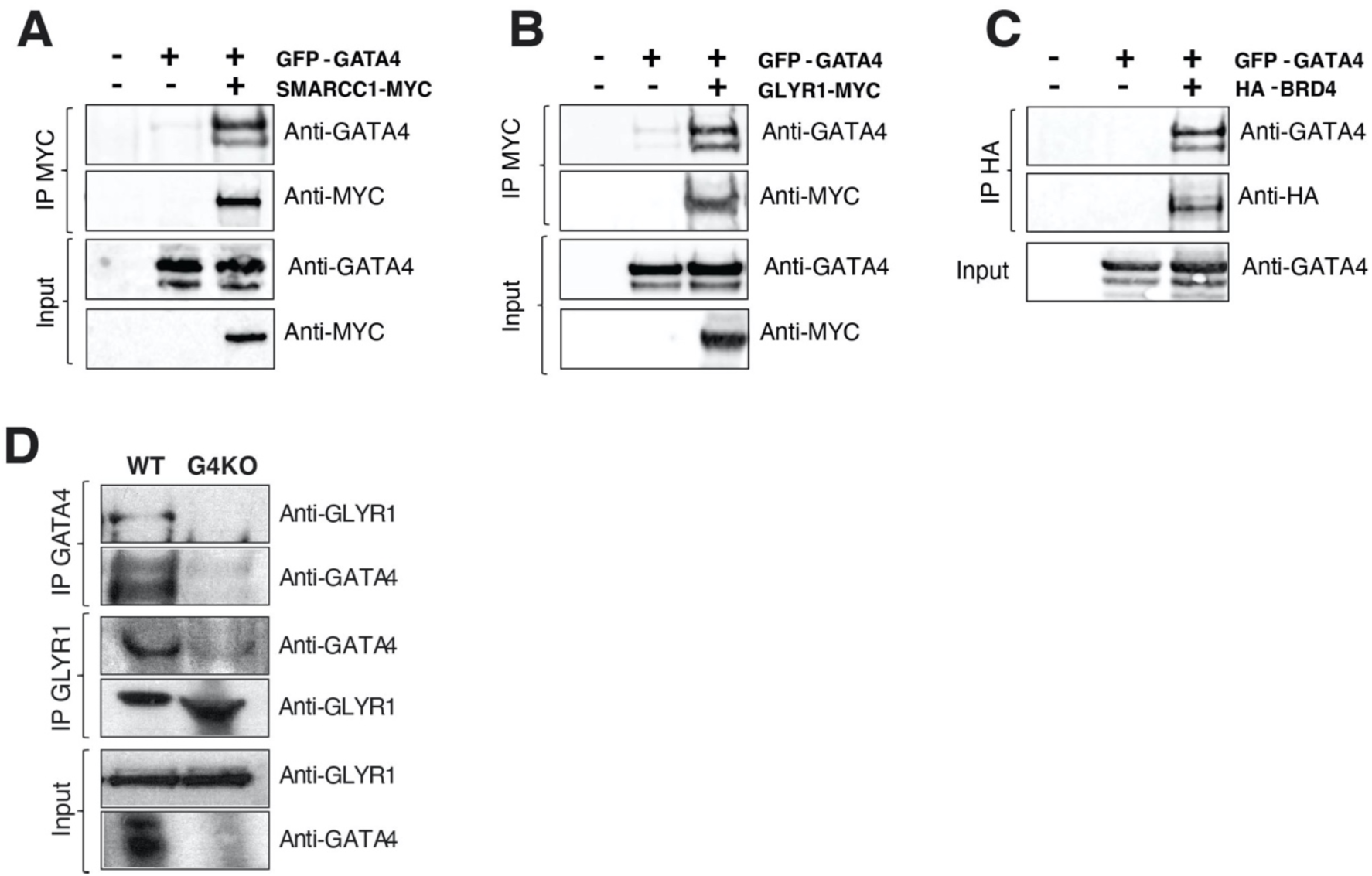
Co-immunoprecipitation validated PPI between top scored CHD candidate genes and GATA4. Related to Figure 4. (A-C) The ability of the proteins encoded by three top-scored interactome CHD candidate genes, SMARCC1 (A), GLYR1 (B) and BRD4 (C), to interact with GATA4 as assessed by ectopic expression of their MYC-or HA-tagged WT proteins in HEK293 cells followed by immunoprecipitation (IP) with anti-MYC or anti-HA antibodies. Enriched nuclear lysates prior to IP (Inputs) were set aside and analyzed by immunoblotting with the indicated antibodies in parallel with IP samples to verify similar protein ectopic expression levels across samples. (D) Immunoprecipitation (IP) for endogenous GATA4 protein and its protein complexes from enriched nuclear lysates of WT and GATA4-KO CPs, followed by immunoblot for indicated antibodies. Aliquots of CP enriched nuclear lysates were put aside prior to IP (Inputs). IP and Inputs were subsequently subjected to immunoblotting with the indicated antibodies.

**Figure S6.**
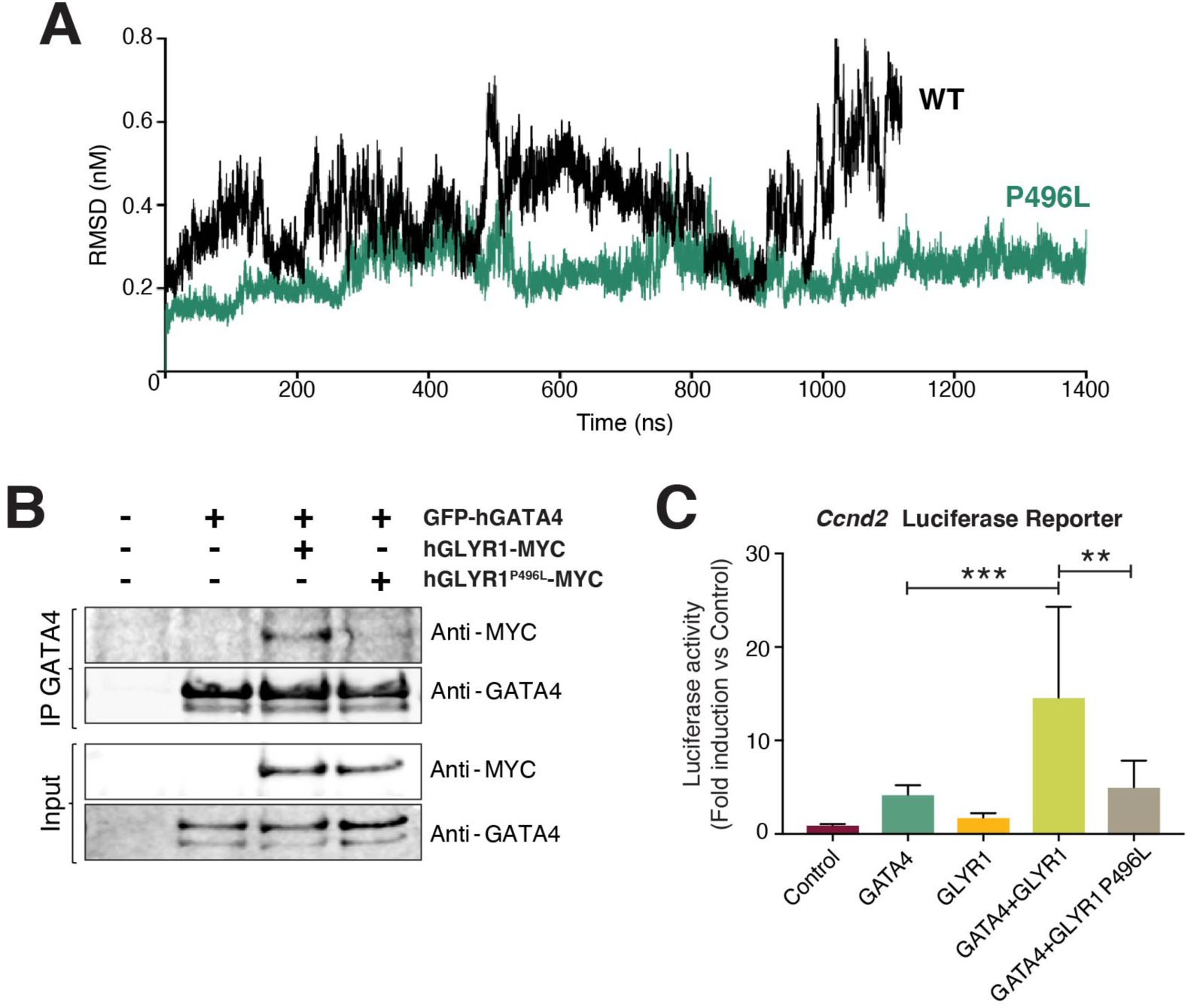
Protein-damaging effect of the CHD missense DNV in GLYR1. Related to Figure 5. (A) Evolution of the root mean square deviation (RMSD) of the structural dynamic frames visited by WT (black) or GLYR1 P496L (green) beta-DH domains over time, taking the starting protein structure as reference. (B) The ability of GLYR1 WT or P496L mutant to interact with GATA4 as assessed by ectopic expression in HEK293 cells and immunoprecipitation (IP) of GFP-GATA4 followed by immunoblotting with the indicated antibodies. Enriched nuclear lysates prior to IP (Inputs) were set aside and analyzed in parallel with IP samples to verify similar protein ectopic expression levels across samples. (C) Luciferase reporter assay in HeLa cells showing activation of the luciferase reporter upon addition of plasmids encoding indicated proteins. Equal amount of total transfected DNA per condition was adjusted with empty vector. (n=3 independent experiments). One-way ANOVA coupled with Tukey post hoc test: **p-value < 0.01, *** p-value <0.001.

**Figure S7.**
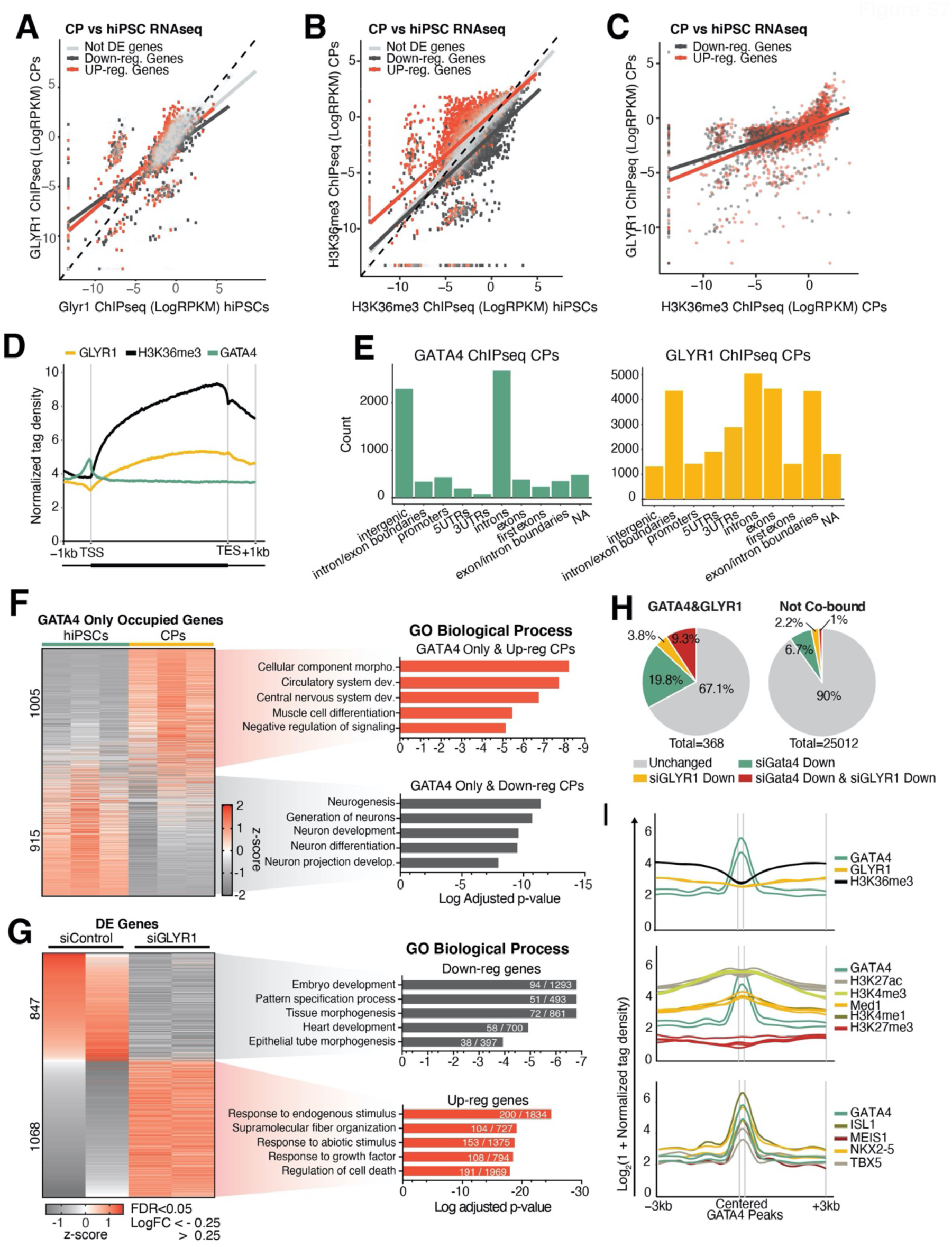
GLYR1 genome-wide occupancy and transcriptional regulation during cardiomyocyte differentiation. Related to Figure 6. (A-C) Scatter plots showing the correlations between indicated ChIPseq signals (log 2 RPKM) at the indicated CP or hiPSC stages for genes classified as not differentially expressed (Not DE genes; light grey), up-regulated (Up-reg genes; red) and down-regulated (Down-reg genes; dark grey) based on publicly available hiPSCs vs. CPs RNAseq data (GSE137920). Dotted lines represent y=x line. ChIPseq GLYR1 hiPSC, H3K36me3 hiPSCs and CPs n=2; GLYR1 CPs ChIPseq n=3. (D) Metagene plot representing the normalized ChIP tag densities for GLYR1, H3K36me3 and GATA4 centered on gene bodies and extending one kilobase upstream of the transcription start sites (TSS) and downstream of the transcription end sites (TES). Curves represent a single representative replicate per ChIP condition. (E) Distribution of GATA4 and GLYR1 genome-wide occupancy across indicated features as assessed by ChIPseq in CPs. GLYR1 CPs ChIPseq n=3; GATA4 ChIPseq n=2. (F) Gene expression and biological functions of genes defined by ChIPseq as occupied by GATA4 only in hiPSC (D0) and CPs (D7). Gene expression is represented as z-score based on publicly available RNAseq data comparing CPs and hiPSCs (GSE137920). Gene Ontology (GO) Biological Process enrichment analysis for GATA4-Only genes up-regulated (Up-reg: CP average z-score>0.1; red) and down-regulated (Down-reg: CP average z-score<-0.1; grey) from hiPSC to CPs. Dev.: development; morpho.: morphogenesis. (G) Genes differentially expressed (DE) upon GLYR1 knockdown at cardiac progenitor stage (FDR<0.05, LogFC<-0.25; n=2). Cells were transfected with Control or GLYR1 siRNAs at day 4 of differentiation and cardiac progenitors collected 72h later for RNAseq. Bar graphs represent enriched Biological Process terms from Gene Ontology (GO) for down-regulated (grey) genes and up-regulated genes (red) in siGLYR1 compared to siControl treated cells. The number of DE genes and the total number of genes in each GO category are indicated in each bar graph. (H) Pie charts showing the percentage of genes differentially expressed (DE; FDR<0.05, LogFC<-0.25) upon GATA4 knockdown (siGATA4), GLYR1 knockdown (siGLYR1), downregulated upon either independently added siRNAs (siGATA4 and siGLYR1), as well as non-DE genes (unchanged) for GATA4&GLYR1-bound genes and Not co-bound genes. siControl vs siGATA4 RNAseq (n=3); siControl vs siGLYR1 RNASeq (n=2). Each replicate corresponds to independent cardiomyocyte differentiations. (I) Metagene plots for GATA4&GLYR1-bound genes showing the cardiac progenitors normalized ChIPseq signal for GATA4 (n=2), GLYR1 (n=2) and H3K36me3 (n=2) (upper panel), the indicated histone marks (middle panel; public available data GSE85631 and GSM2047027) and the GATA4 (n=2), TBX5 (n=2), NKX2-5 (n=2), MEIS1 (n=1) and ISL1 (n=1) cTFs centered on GATA4 peaks inside the gene body (1^st^ Intron-TES).

**Table S1. WT versus GATA4-KO CPs differential mRNA expression analysis. Related to** Figures 1 **and** S1.

**Table S2. WT versus TBX5-KO CPs differential mRNA expression analysis. Related to** Figures 1 **and** S1.

**Table S3. APMS data and filtering criteria for the GATA4 and TBX5 interactome in CPs. Related to** Figures 1 **and** S2.

**Table S4. Permutation-based statistical analysis of GT-PPIs enrichment in CHD-associated genomic variants. Related to** Figure 2.

**Table S5. Proband variants in genes involved in mouse/human heart development (Jin et al., 2017) removed from the CP interactome (Heart Dev. Unknown) Permutation analysis in** Figure 2B**. Related to** Figure 2.

**Table S6: APMS data and filtering criteria for the GATA4 and TBX5 interactome in HEK293T cells. Related to** Figure 2 **and** S3.

**Table S7: De novo missense, loss-of-function and splice PCGC variants found in GT-PPI genes. Related to** Figure 3 **and** S4.

**Table S8. Counts for the different types of de novo variants (DNVs) found in CHD and control cohorts in GT-PPIs and CHD variant network enrichment analysis.**

**Table S9. Percentile of expression in the developing heart for interactome genes with reported protein-altering DNVs found in the CHD cohort. Related to** Figure 4.

**Table S10. Integrative pathogenesis scoring of interactome protein-altering missense DNVs found in CHD patients. Related to** Figure 4.

**Table S11. Table for GLYR1 differentially bound genes between hiPSCs and CPs subjected to k-means clustering based on GLYR1 ChIPseq signal, H3K36me3 ChIPseq signal and gene expression levels. Related to** Figure 6.

**Table S12. GO enrichment analysis for Clusters 1,2 and 3 in** Figure 6A**. Related to** Figure 6.

**Table S13. List of GATA4 & GLYR1, GATA4-only or GLYR1-only bound genes within the gene body (1st Intron-TES). Related to** Figure 6.

**Table S14. GO enrichment analysis for GATA4 & GLYR1, GATA4-only or GLYR1-only bound genes within the gene body (1st Intron-TES). Related to** Figure 6 **and** S7.

**Table S15: GATA4-only bound gene expression in hiPSCs and cardiac progenitors and z-score calculation based on GSE137920 RNAseq;** Figure S7F

**Table S16. SiControl versus siGATA4 differential mRNA expression analysis in hiPSC-derived cardiac progenitors. Related to** Figure 6.

**Table S17. SiControl versus siGLYR1 differential mRNA expression analysis in hiPSC-derived cardiac progenitors. Related to** Figure 6 **and** S7.

**Table S18. GO term enrichment analysis for DE genes in siGLYR1 vs siControl cardiac progenitors. Related to** Figure S7.

## REFERENCES

Akerberg, B.N., Gu, F., VanDusen, N.J., Zhang, X., Dong, R., Li, K., Zhang, B., Zhou, B., Sethi, I., Ma, Q., et al. (2019). A reference map of murine cardiac transcription factor chromatin occupancy identifies dynamic and conserved enhancers. Nat. Commun. 10, 4907.

Andrews, S. (2007). A quality control tool for high throughput sequence data. (babraham bioinformatics).

Ang, Y.-S., Rivas, R.N., Ribeiro, A.J.S., Srivas, R., Rivera, J., Stone, N.R., Pratt, K., Mohamed, T.M.A., Fu, J.-D., Spencer, C.I., et al. (2016). Disease model of GATA4 mutation reveals transcription factor cooperativity in human cardiogenesis. Cell 167, 1734–1749.e22.

Aronesty, E. (2013). Comparison of Sequencing Utility Programs. Open Bioinforma. J. 7, 1–8.

Baardman, M.E., Zwier, M.V., Wisse, L.J., Gittenberger-de Groot, A.C., Kerstjens-Frederikse, W.S., Hofstra, R.M.W., Jurdzinski, A., Hierck, B.P., Jongbloed, M.R.M., Berger, R.M.F., et al. (2016). Common arterial trunk and ventricular non-compaction in Lrp2 knockout mice indicate a crucial role of LRP2 in cardiac development. Dis. Model. Mech. 9, 413–425.

Barrett, T., Wilhite, S.E., Ledoux, P., Evangelista, C., Kim, I.F., Tomashevsky, M., Marshall, K.A., Phillippy, K.H., Sherman, P.M., Holko, M., et al. (2013). NCBI GEO: archive for functional genomics data sets--update. Nucleic Acids Res. 41, D991–5.

Basson, C.T., Bachinsky, D.R., Lin, R.C., Levi, T., Elkins, J.A., Soults, J., Grayzel, D., Kroumpouzou, E., Traill, T.A., Leblanc-Straceski, J., et al. (1997). Mutations in human TBX5 [corrected] cause limb and cardiac malformation in Holt-Oram syndrome. Nat. Genet. 15, 30–35.

Basson, C.T., Huang, T., Lin, R.C., Bachinsky, D.R., Weremowicz, S., Vaglio, A., Bruzzone, R., Quadrelli, R., Lerone, M., Romeo, G., et al. (1999). Different TBX5 interactions in heart and limb defined by Holt-Oram syndrome mutations. Proc. Natl. Acad. Sci. USA 96, 2919–2924.

Bekker, H., Berendsen, H., Dijkstra, E., Achterop, S., van Drunen, R., Van Der Spoel, D., Sijbers, A., Keegstra, H., Reitsma, B., Renardus, M., et al. (1993). Gromacs: A parallel computer for molecular dynamics simulations – ScienceOpen.

Bouman, A., Alders, M., Oostra, R.J., van Leeuwen, E., Thuijs, N., van der Kevie-Kersemaekers, A.-M., and van Maarle, M. (2017). Oral-facial-digital syndrome type 1 in males: Congenital heart defects are included in its phenotypic spectrum. Am. J. Med. Genet. A 173, 1383–1389.

Bruneau, B.G., Logan, M., Davis, N., Levi, T., Tabin, C.J., Seidman, J.G., and Seidman, C.E. (1999). Chamber-specific cardiac expression of Tbx5 and heart defects in Holt-Oram syndrome. Dev. Biol. 211, 100–108.

Bruneau, B.G., Nemer, G., Schmitt, J.P., Charron, F., Robitaille, L., Caron, S., Conner, D.A., Gessler, M., Nemer, M., Seidman, C.E., et al. (2001). A murine model of Holt-Oram syndrome defines roles of the T-box transcription factor Tbx5 in cardiogenesis and disease. Cell 106, 709– 721.

Bryois, J., Skene, N.G., Hansen, T.F., Kogelman, L.J.A., Watson, H.J., Liu, Z., Eating Disorders Working Group of the Psychiatric Genomics Consortium, International Headache Genetics Consortium, 23andMe Research Team, Brueggeman, L., et al. (2020). Genetic identification of cell types underlying brain complex traits yields insights into the etiology of Parkinson’s disease. Nat. Genet. 52, 482–493.

Caldera, M., Buphamalai, P., Müller, F., and Menche, J. (2017). Interactome-based approaches to human disease. Current Opinion in Systems Biology 3, 88–94.

Camenisch, T.D., Spicer, A.P., Brehm-Gibson, T., Biesterfeldt, J., Augustine, M.L., Calabro, A., Kubalak, S., Klewer, S.E., and McDonald, J.A. (2000). Disruption of hyaluronan synthase-2 abrogates normal cardiac morphogenesis and hyaluronan-mediated transformation of epithelium to mesenchyme. J. Clin. Invest. 106, 349–360.

Case, D.A., Cerutti, D.S., Cheatham, T.E.I., Darden, T.A., Duke, R.E., Giese, T.J., Gohlke, H., Goetz, A.W., Greene, D., Homeyer, N., et al. (2017). Amber18 (University of San Francisco).

Castillo-Robles, J., Ramírez, L., Spaink, H.P., and Lomelí, H. (2018). smarce1 mutants have a defective endocardium and an increased expression of cardiac transcription factors in zebrafish. Sci. Rep. 8, 15369.

Cavalcante, R.G., and Sartor, M.A. (2017). annotatr: genomic regions in context. Bioinformatics 33, 2381–2383.

Chen, J., Yuan, H., Xie, K., Wang, X., Tan, L., Zou, Y., Yang, Y., Pan, L., Xiao, J., Chen, G., et al. (2020). A novel TAB2 nonsense mutation (p.S149X) causing autosomal dominant congenital heart defects: a case report of a Chinese family. BMC Cardiovasc. Disord. 20, 27.

Christianson, A., and Howson, C.P. (2006). March of dimes. Global Report on Birth.

Cox, J., and Mann, M. (2008). MaxQuant enables high peptide identification rates, individualized p.p.b.-range mass accuracies and proteome-wide protein quantification. Nat. Biotechnol. 26, 1367–1372.

Darden, T., York, D., and Pedersen, L. (1993). Particle mesh Ewald: An N⋅log(N) method for Ewald sums in large systems. J. Chem. Phys. 98, 10089.

Deciphering Developmental Disorders Study (2015). Large-scale discovery of novel genetic causes of developmental disorders. Nature 519, 223–228.

Diets, I.J., Prescott, T., Champaigne, N.L., Mancini, G.M.S., Krossnes, B., Frič, R., Kocsis, K., Jongmans, M.C.J., and Kleefstra, T. (2019). A recurrent de novo missense pathogenic variant in SMARCB1 causes severe intellectual disability and choroid plexus hyperplasia with resultant hydrocephalus. Genet. Med. 21, 572–579.

Dobin, A., Davis, C.A., Schlesinger, F., Drenkow, J., Zaleski, C., Jha, S., Batut, P., Chaisson, M., and Gingeras, T.R. (2013). STAR: ultrafast universal RNA-seq aligner. Bioinformatics 29, 15–21.

Dsouza, N.R., Zimmermann, M.T., and Geddes, G.C. (2019). A case of Coffin-Siris syndrome with severe congenital heart disease and a novel SMARCA4 variant. Cold Spring Harb Mol Case Stud 5.

Dupays, L., Shang, C., Wilson, R., Kotecha, S., Wood, S., Towers, N., and Mohun, T. (2015). Sequential Binding of MEIS1 and NKX2-5 on the Popdc2 Gene: A Mechanism for Spatiotemporal Regulation of Enhancers during Cardiogenesis. Cell Rep. 13, 183–195.

Enane, F.O., Shuen, W.H., Gu, X., Quteba, E., Przychodzen, B., Makishima, H., Bodo, J., Ng, J., Chee, C.L., Ba, R., et al. (2017). GATA4 loss of function in liver cancer impedes precursor to hepatocyte transition. J. Clin. Invest. 127, 3527–3542.

Fang, R., Chen, F., Dong, Z., Hu, D., Barbera, A.J., Clark, E.A., Fang, J., Yang, Y., Mei, P., Rutenberg, M., et al. (2013). LSD2/KDM1B and its cofactor NPAC/GLYR1 endow a structural and molecular model for regulation of H3K4 demethylation. Mol. Cell 49, 558–570.

Fei, J., Ishii, H., Hoeksema, M.A., Meitinger, F., Kassavetis, G.A., Glass, C.K., Ren, B., and Kadonaga, J.T. (2018). NDF, a nucleosome-destabilizing factor that facilitates transcription through nucleosomes. Genes Dev. 32, 682–694.

Ferrante, M.I., Zullo, A., Barra, A., Bimonte, S., Messaddeq, N., Studer, M., Dollé, P., and Franco, B. (2006). Oral-facial-digital type I protein is required for primary cilia formation and left-right axis specification. Nat. Genet. 38, 112–117.

Fu, J., Yang, Z., Wei, J., Han, J., and Gu, J. (2006). Nuclear protein NP60 regulates p38 MAPK activity. J. Cell Sci. 119, 115–123.

Fuller, Z.L., Berg, J.J., Mostafavi, H., Sella, G., and Przeworski, M. (2019). Measuring intolerance to mutation in human genetics. Nat. Genet. 51, 772–776.

Furtado, M.B., Wilmanns, J.C., Chandran, A., Perera, J., Hon, O., Biben, C., Willow, T.J., Nim, H.T., Kaur, G., Simonds, S., et al. (2017). Point mutations in murine Nkx2-5 phenocopy human congenital heart disease and induce pathogenic Wnt signaling. JCI Insight 2, e88271.

Garg, V., Kathiriya, I.S., Barnes, R., Schluterman, M.K., King, I.N., Butler, C.A., Rothrock, C.R., Eapen, R.S., Hirayama-Yamada, K., Joo, K., et al. (2003). GATA4 mutations cause human congenital heart defects and reveal an interaction with TBX5. Nature 424, 443–447.

Gifford, C.A., Ranade, S.S., Samarakoon, R., Salunga, H.T., de Soysa, T.Y., Huang, Y., Zhou, P., Elfenbein, A., Wyman, S.K., Bui, Y.K., et al. (2019). Oligogenic inheritance of a human heart disease involving a genetic modifier. Science 364, 865–870.

Goh, K.-I., Cusick, M.E., Valle, D., Childs, B., Vidal, M., and Barabási, A.-L. (2007). The human disease network. Proc. Natl. Acad. Sci. USA 104, 8685–8690.

González-Terán, B., López, J.A., Rodríguez, E., Leiva, L., Martínez-Martínez, S., Bernal, J.A., Jiménez-Borreguero, L.J., Redondo, J.M., Vazquez, J., and Sabio, G. (2016). p38γ and δ promote heart hypertrophy by targeting the mTOR-inhibitory protein DEPTOR for degradation. Nat. Commun. 7, 10477.

Gordillo, M., Vega, H., and Jabs, E.W. (1993). Roberts Syndrome. In GeneReviews(®), R.A. Pagon, M.P. Adam, H.H. Ardinger, S.E. Wallace, A. Amemiya, L.J. Bean, T.D. Bird, N. Ledbetter, H.C. Mefford, R.J. Smith, et al., eds. (Seattle (WA): University of Washington, Seattle), p.

GTEx Consortium, Laboratory, Data Analysis &Coordinating Center (LDACC)—Analysis Working Group, Statistical Methods groups—Analysis Working Group, Enhancing GTEx (eGTEx) groups, NIH Common Fund, NIH/NCI, NIH/NHGRI, NIH/NIMH, NIH/NIDA, Biospecimen Collection Source Site—NDRI, et al. (2017). Genetic effects on gene expression across human tissues. Nature 550, 204–213.

Guo, Y., Mahony, S., and Gifford, D.K. (2012). High resolution genome wide binding event finding and motif discovery reveals transcription factor spatial binding constraints. PLoS Comput. Biol. 8, e1002638.

Hekselman, I., and Yeger-Lotem, E. (2020). Mechanisms of tissue and cell-type specificity in heritable traits and diseases. Nat. Rev. Genet. 21, 137–150.

Heyne, H.O., Singh, T., Stamberger, H., Abou Jamra, R., Caglayan, H., Craiu, D., De Jonghe, P., Guerrini, R., Helbig, K.L., Koeleman, B.P.C., et al. (2018). De novo variants in neurodevelopmental disorders with epilepsy. Nat. Genet. 50, 1048–1053.

Hinton, R.B., Prakash, A., Romp, R.L., Krueger, D.A., Knilans, T.K., and International Tuberous Sclerosis Consensus Group (2014). Cardiovascular manifestations of tuberous sclerosis complex and summary of the revised diagnostic criteria and surveillance and management recommendations from the International Tuberous Sclerosis Consensus Group. J. Am. Heart Assoc. 3, e001493.

Homsy, J., Zaidi, S., Shen, Y., Ware, J.S., Samocha, K.E., Karczewski, K.J., DePalma, S.R., McKean, D., Wakimoto, H., Gorham, J., et al. (2015). De novo mutations in congenital heart disease with neurodevelopmental and other congenital anomalies. Science 350, 1262–1266.

Hota, S.K., and Bruneau, B.G. (2016). ATP-dependent chromatin remodeling during mammalian development. Development 143, 2882–2897.

Hu, X., Li, T., Zhang, C., Liu, Y., Xu, M., Wang, W., Jia, Z., Ma, K., Zhang, Y., and Zhou, C. (2011). GATA4 regulates ANF expression synergistically with Sp1 in a cardiac hypertrophy model. J. Cell Mol. Med. 15, 1865–1877.

Huang, N., Lee, I., Marcotte, E.M., and Hurles, M.E. (2010). Characterising and predicting haploinsufficiency in the human genome. PLoS Genet. 6, e1001154.

Izarzugaza, J.M.G., Ellesøe, S.G., Doganli, C., Ehlers, N.S., Dalgaard, M.D., Audain, E., Dombrowsky, G., Sifrim, A., Wilsdon, A., Thienpont, B., et al. (2019). Systems genetics analysis identify calcium signalling defects as novel cause of congenital heart disease. BioRxiv.

Ji, W., Ferdman, D., Copel, J., Scheinost, D., Shabanova, V., Brueckner, M., Khokha, M.K., and Ment, L.R. (2020). De novo damaging variants associated with congenital heart diseases contribute to the connectome. Sci. Rep. 10, 7046.

Jimenez-Morales, D., Rosa Campos, A., Von Dollen, J., and Swaney, D. (2020). artMS: Analytical R tools for Mass Spectrometry version 1.6.5 from Bioconductor (Bioconductor).

Jin, S.C., Homsy, J., Zaidi, S., Lu, Q., Morton, S., DePalma, S.R., Zeng, X., Qi, H., Chang, W., Sierant, M.C., et al. (2017). Contribution of rare inherited and de novo variants in 2,871 congenital heart disease probands. Nat. Genet. 49, 1593–1601.

Jones, W.D., Dafou, D., McEntagart, M., Woollard, W.J., Elmslie, F.V., Holder-Espinasse, M., Irving, M., Saggar, A.K., Smithson, S., Trembath, R.C., et al. (2012). De novo mutations in MLL cause Wiedemann-Steiner syndrome. Am. J. Hum. Genet. 91, 358–364.

Jorgensen, W.L., Chandrasekhar, J., Madura, J.D., Impey, R.W., and Klein, M.L. (1983). Comparison of simple potential functions for simulating liquid water. J. Chem. Phys. 79, 926.

Karczewski, K.J., Francioli, L.C., Tiao, G., Cummings, B.B., Alföldi, J., Wang, Q., Collins, R.L., Laricchia, K.M., Ganna, A., Birnbaum, D.P., et al. (2020). The mutational constraint spectrum quantified from variation in 141,456 humans. Nature 581, 434–443.

Kimura, H. (2013). Histone modifications for human epigenome analysis. J. Hum. Genet. 58, 439– 445.

Knowlton, K.U., Baracchini, E., Ross, R.S., Harris, A.N., Henderson, S.A., Evans, S.M., Glembotski, C.C., and Chien, K.R. (1991). Co-regulation of the atrial natriuretic factor and cardiac myosin light chain-2 genes during alpha-adrenergic stimulation of neonatal rat ventricular cells. Identification of cis sequences within an embryonic and a constitutive contractile protein gene which mediate inducible expression. J. Biol. Chem. 266, 7759–7768.

Kuo, C.T., Morrisey, E.E., Anandappa, R., Sigrist, K., Lu, M.M., Parmacek, M.S., Soudais, C., and Leiden, J.M. (1997). GATA4 transcription factor is required for ventral morphogenesis and heart tube formation. Genes Dev. 11, 1048–1060.

Lambert, S.A., Jolma, A., Campitelli, L.F., Das, P.K., Yin, Y., Albu, M., Chen, X., Taipale, J., Hughes, T.R., and Weirauch, M.T. (2018). The human transcription factors. Cell 172, 650–665.

Langmead, B., and Salzberg, S.L. (2012). Fast gapped-read alignment with Bowtie 2. Nat. Methods 9, 357–359.

Lau, E., Han, Y., Williams, D.R., Thomas, C.T., Shrestha, R., Wu, J.C., and Lam, M.P.Y. (2019). Splice-Junction-Based Mapping of Alternative Isoforms in the Human Proteome. Cell Rep. 29, 3751–3765.e5.

Lebrun, N., Giurgea, I., Goldenberg, A., Dieux, A., Afenjar, A., Ghoumid, J., Diebold, B., Mietton, L., Briand-Suleau, A., Billuart, P., et al. (2018). Molecular and cellular issues of KMT2A variants involved in Wiedemann-Steiner syndrome. Eur. J. Hum. Genet. 26, 107–116.

Lei, I., Gao, X., Sham, M.H., and Wang, Z. (2012). SWI/SNF protein component BAF250a regulates cardiac progenitor cell differentiation by modulating chromatin accessibility during second heart field development. J. Biol. Chem. 287, 24255–24262.

Lepore, J.J., Mericko, P.A., Cheng, L., Lu, M.M., Morrisey, E.E., and Parmacek, M.S. (2006). GATA-6 regulates semaphorin 3C and is required in cardiac neural crest for cardiovascular morphogenesis. J. Clin. Invest. 116, 929–939.

Li, Q.Y., Newbury-Ecob, R.A., Terrett, J.A., Wilson, D.I., Curtis, A.R., Yi, C.H., Gebuhr, T., Bullen, P.J., Robson, S.C., Strachan, T., et al. (1997). Holt-Oram syndrome is caused by mutations in TBX5, a member of the Brachyury (T) gene family. Nat. Genet. 15, 21–29.

Lian, X., Zhang, J., Azarin, S.M., Zhu, K., Hazeltine, L.B., Bao, X., Hsiao, C., Kamp, T.J., and Palecek, S.P. (2013). Directed cardiomyocyte differentiation from human pluripotent stem cells by modulating Wnt/β-catenin signaling under fully defined conditions. Nat. Protoc. 8, 162–175.

Liao, Y., Smyth, G.K., and Shi, W. (2014). featureCounts: an efficient general purpose program for assigning sequence reads to genomic features. Bioinformatics 30, 923–930.

Lun, A.T.L., Chen, Y., and Smyth, G.K. (2016). It’s DE-licious: A Recipe for Differential Expression Analyses of RNA-seq Experiments Using Quasi-Likelihood Methods in edgeR. Methods Mol. Biol. 1418, 391–416.

Luna-Zurita, L., Stirnimann, C.U., Glatt, S., Kaynak, B.L., Thomas, S., Baudin, F., Samee, M.A.H., He, D., Small, E.M., Mileikovsky, M., et al. (2016). Complex interdependence regulates heterotypic transcription factor distribution and coordinates cardiogenesis. Cell 164, 999–1014.

Maere, S., Heymans, K., and Kuiper, M. (2005). BiNGO: a Cytoscape plugin to assess overrepresentation of gene ontology categories in biological networks. Bioinformatics 21, 3448– 3449.

Maestro Schrödinger, LLC (2019). Maestro Suite of Programs (v. 2019−4) (Maestro Schrödinger, LLC).

Magger, O., Waldman, Y.Y., Ruppin, E., and Sharan, R. (2012). Enhancing the prioritization of disease-causing genes through tissue specific protein interaction networks. PLoS Comput. Biol. 8, e1002690.

Maitra, M., Schluterman, M.K., Nichols, H.A., Richardson, J.A., Lo, C.W., Srivastava, D., and Garg, V. (2009). Interaction of Gata4 and Gata6 with Tbx5 is critical for normal cardiac development. Dev. Biol. 326, 368–377.

Maitra, M., Koenig, S.N., Srivastava, D., and Garg, V. (2010). Identification of GATA6 sequence variants in patients with congenital heart defects. Pediatr. Res. 68, 281–285.

Marabelli, C., Marrocco, B., Pilotto, S., Chittori, S., Picaud, S., Marchese, S., Ciossani, G., Forneris, F., Filippakopoulos, P., Schoehn, G., et al. (2019). A Tail-Based Mechanism Drives Nucleosome Demethylation by the LSD2/NPAC Multimeric Complex. Cell Rep. 27, 387–399.e7.

Miyamoto, S., and Kollman, P.A. (1992). Settle: An analytical version of the SHAKE and RATTLE algorithm for rigid water models. J. Comput. Chem. 13, 952–962.

Miyaoka, Y., Chan, A.H., Judge, L.M., Yoo, J., Huang, M., Nguyen, T.D., Lizarraga, P.P., So, P.-L., and Conklin, B.R. (2014). Isolation of single-base genome-edited human iPS cells without antibiotic selection. Nat. Methods 11, 291–293.

Molkentin, J.D., Lin, Q., Duncan, S.A., and Olson, E.N. (1997). Requirement of the transcription factor GATA4 for heart tube formation and ventral morphogenesis. Genes Dev. 11, 1061–1072.

Montefiori, M., Pilotto, S., Marabelli, C., Moroni, E., Ferraro, M., Serapian, S.A., Mattevi, A., and Colombo, G. (2019). Impact of Mutations on NPAC Structural Dynamics: Mechanistic Insights from MD Simulations. J. Chem. Inf. Model. 59, 3927–3937.

Mori, A.D., Zhu, Y., Vahora, I., Nieman, B., Koshiba-Takeuchi, K., Davidson, L., Pizard, A., Seidman, J.G., Seidman, C.E., Chen, X.J., et al. (2006). Tbx5-dependent rheostatic control of cardiac gene expression and morphogenesis. Dev. Biol. 297, 566–586.

Moskowitz, I.P., Wang, J., Peterson, M.A., Pu, W.T., Mackinnon, A.C., Oxburgh, L., Chu, G.C., Sarkar, M., Berul, C., Smoot, L., et al. (2011). Transcription factor genes Smad4 and Gata4 cooperatively regulate cardiac valve development. [corrected]. Proc. Natl. Acad. Sci. USA 108, 4006–4011.

Nakamura, H., Cook, R.N., and Justice, M.J. (2013). Mouse Tenm4 is required for mesoderm induction. BMC Dev. Biol. 13, 9.

Narita, N., Bielinska, M., and Wilson, D.B. (1997). Cardiomyocyte differentiation by GATA-4-deficient embryonic stem cells. Development 124, 3755–3764.

Padmanabhan, A., Alexanian, M., Linares-Saldana, R., González-Terán, B., Andreoletti, G., Huang, Y., Connolly, A.J., Kim, W., Hsu, A., Duan, Q., et al. (2020). BRD4 Interacts with GATA4 to Govern Mitochondrial Homeostasis in Adult Cardiomyocytes. Circulation.

Parisot, P., Bajolle, F., Attié-Bittach, T., Thomas, S., Goudefroye, G., Abadie, V., Lyonnet, S., and Bonnet, D. (2010). 321 Congenital heart defects in CHARGE syndrome patients with CHD7 mutations. Archives of Cardiovascular Diseases Supplements 2, 104–105.

Park, C.Y., Pierce, S.A., von Drehle, M., Ivey, K.N., Morgan, J.A., Blau, H.M., and Srivastava, D. (2010). skNAC, a Smyd1-interacting transcription factor, is involved in cardiac development and skeletal muscle growth and regeneration. Proc. Natl. Acad. Sci. USA 107, 20750–20755.

Perez-Riverol, Y., Csordas, A., Bai, J., Bernal-Llinares, M., Hewapathirana, S., Kundu, D.J., Inuganti, A., Griss, J., Mayer, G., Eisenacher, M., et al. (2019). The PRIDE database and related tools and resources in 2019: improving support for quantification data. Nucleic Acids Res. 47, D442–D450.

Pierpont, M.E., Brueckner, M., Chung, W.K., Garg, V., Lacro, R.V., McGuire, A.L., Mital, S., Priest, J.R., Pu, W.T., Roberts, A., et al. (2018). Genetic basis for congenital heart disease: revisited: A scientific statement from the american heart association. Circulation 138, e653–e711.

R Core Team (2020). R: A language and environment for statistical computing. (R Foundation for Statistical Computing, Vienna, Austria.).

Rahman, S., Sowa, M.E., Ottinger, M., Smith, J.A., Shi, Y., Harper, J.W., and Howley, P.M. (2011). The Brd4 extraterminal domain confers transcription activation independent of pTEFb by recruiting multiple proteins, including NSD3. Mol. Cell. Biol. 31, 2641–2652.

Rajagopal, S.K., Ma, Q., Obler, D., Shen, J., Manichaikul, A., Tomita-Mitchell, A., Boardman, K., Briggs, C., Garg, V., Srivastava, D., et al. (2007). Spectrum of heart disease associated with murine and human GATA4 mutation. J. Mol. Cell Cardiol. 43, 677–685.

Ramírez, F., Ryan, D.P., Grüning, B., Bhardwaj, V., Kilpert, F., Richter, A.S., Heyne, S., Dündar, F., and Manke, T. (2016). deepTools2: a next generation web server for deep-sequencing data analysis. Nucleic Acids Res. 44, W160–5.

Rasmussen, T.L., Ma, Y., Park, C.Y., Harriss, J., Pierce, S.A., Dekker, J.D., Valenzuela, N., Srivastava, D., Schwartz, R.J., Stewart, M.D., et al. (2015). Smyd1 facilitates heart development by antagonizing oxidative and ER stress responses. PLoS One 10, e0121765.

Razick, S., Magklaras, G., and Donaldson, I.M. (2008). iRefIndex: a consolidated protein interaction database with provenance. BMC Bioinformatics 9, 405.

Rentzsch, P., Witten, D., Cooper, G.M., Shendure, J., and Kircher, M. (2019). CADD: predicting the deleteriousness of variants throughout the human genome. Nucleic Acids Res. 47, D886–D894.

Richter, F., Morton, S.U., Kim, S.W., Kitaygorodsky, A., Wasson, L.K., Chen, K.M., Zhou, J., Qi, H., Patel, N., DePalma, S.R., et al. (2020). Genomic analyses implicate noncoding de novo variants in congenital heart disease. Nat. Genet. 52, 769–777.

Robinson, M.D., and Oshlack, A. (2010). A scaling normalization method for differential expression analysis of RNA-seq data. Genome Biol. 11, R25.

Robinson, M.D., and Smyth, G.K. (2007). Moderated statistical tests for assessing differences in tag abundance. Bioinformatics 23, 2881–2887.

Robinson, M.D., and Smyth, G.K. (2008). Small-sample estimation of negative binomial dispersion, with applications to SAGE data. Biostatistics 9, 321–332.

Robinson, M.D., McCarthy, D.J., and Smyth, G.K. (2010). edgeR: a Bioconductor package for differential expression analysis of digital gene expression data. Bioinformatics 26, 139–140.

Saito, Y., Kojima, T., and Takahashi, N. (2012). Mab21l2 is essential for embryonic heart and liver development. PLoS One 7, e32991.

Sastry, G.M., Adzhigirey, M., Day, T., Annabhimoju, R., and Sherman, W. (2013). Protein and ligand preparation: parameters, protocols, and influence on virtual screening enrichments. J. Comput. Aided Mol. Des. 27, 221–234.

Shannon, P., Markiel, A., Ozier, O., Baliga, N.S., Wang, J.T., Ramage, D., Amin, N., Schwikowski, B., and Ideker, T. (2003). Cytoscape: a software environment for integrated models of biomolecular interaction networks. Genome Res. 13, 2498–2504.

Sifrim, A., Hitz, M.-P., Wilsdon, A., Breckpot, J., Turki, S.H.A., Thienpont, B., McRae, J., Fitzgerald, T.W., Singh, T., Swaminathan, G.J., et al. (2016). Distinct genetic architectures for syndromic and nonsyndromic congenital heart defects identified by exome sequencing. Nat. Genet. 48, 1060–1065.

Smyth, G.K. (1996). A conditional approach to residual maximum likelihood estimation in generalized linear models (R. Stat. Soc. B.).

de Soysa, T.Y., Ranade, S.S., Okawa, S., Ravichandran, S., Huang, Y., Salunga, H.T., Schricker, A., Del Sol, A., Gifford, C.A., and Srivastava, D. (2019). Single-cell analysis of cardiogenesis reveals basis for organ-level developmental defects. Nature 572, 120–124.

Takeuchi, J.K., Lou, X., Alexander, J.M., Sugizaki, H., Delgado-Olguín, P., Holloway, A.K., Mori, A.D., Wylie, J.N., Munson, C., Zhu, Y., et al. (2011). Chromatin remodelling complex dosage modulates transcription factor function in heart development. Nat. Commun. 2, 187.

Teo, G., Koh, H., Fermin, D., Lambert, J.-P., Knight, J.D.R., Gingras, A.-C., and Choi, H. (2016). SAINTq: Scoring protein-protein interactions in affinity purification - mass spectrometry experiments with fragment or peptide intensity data. Proteomics 16, 2238–2245.

Theis, J.L., Vogler, G., Missinato, M.A., Li, X., Martinez-Fernandez, A., Nielsen, T., Walls, S.M., Kervadec, A., Zeng, X.-X.I., Kezos, J.N., et al. (2019). Patient-specific functional genomics and disease modeling suggest a role for LRP2 in hypoplastic left heart syndrome. BioRxiv.

Thienpont, B., Zhang, L., Postma, A.V., Breckpot, J., Tranchevent, L.-C., Van Loo, P., Møllgård, K., Tommerup, N., Bache, I., Tümer, Z., et al. (2010). Haploinsufficiency of TAB2 causes congenital heart defects in humans. Am. J. Hum. Genet. 86, 839–849.

Tickle, J., Pilka, E.S., Bunkoczi, G., Berridge, G., Smee, C, Kavanagh, K.L., Hozjan, V., Niesen, F.H, Papagrigoriou, E., Pike, A.C.W., et al. (2007). Structure of the cytokine-like nuclear factor n-pac.

Tohyama, S., Hattori, F., Sano, M., Hishiki, T., Nagahata, Y., Matsuura, T., Hashimoto, H., Suzuki, T., Yamashita, H., Satoh, Y., et al. (2013). Distinct metabolic flow enables large-scale purification of mouse and human pluripotent stem cell-derived cardiomyocytes. Cell Stem Cell 12, 127–137.

Tomita-Mitchell, A., Maslen, C.L., Morris, C.D., Garg, V., and Goldmuntz, E. (2007). GATA4 sequence variants in patients with congenital heart disease. J. Med. Genet. 44, 779–783.

Uhlén, M., Fagerberg, L., Hallström, B.M., Lindskog, C., Oksvold, P., Mardinoglu, A., Sivertsson, Å., Kampf, C., Sjöstedt, E., Asplund, A., et al. (2015). Proteomics. Tissue-based map of the human proteome. Science 347, 1260419.

Van Dijck, A., Vulto-van Silfhout, A.T., Cappuyns, E., van der Werf, I.M., Mancini, G.M., Tzschach, A., Bernier, R., Gozes, I., Eichler, E.E., Romano, C., et al. (2019). Clinical presentation of a complex neurodevelopmental disorder caused by mutations in *ADNP*. Biol. Psychiatry 85, 287–297.

Waldron, L., Steimle, J.D., Greco, T.M., Gomez, N.C., Dorr, K.M., Kweon, J., Temple, B., Yang, X.H., Wilczewski, C.M., Davis, I.J., et al. (2016). The cardiac TBX5 interactome reveals a chromatin remodeling network essential for cardiac septation. Dev. Cell 36, 262–275.

Wang, L., Wang, S., and Li, W. (2012). RSeQC: quality control of RNA-seq experiments. Bioinformatics 28, 2184–2185.

Wilczewski, C.M., Hepperla, A.J., Shimbo, T., Wasson, L., Robbe, Z.L., Davis, I.J., Wade, P.A., and Conlon, F.L. (2018). CHD4 and the NuRD complex directly control cardiac sarcomere formation. Proc. Natl. Acad. Sci. USA 115, 6727–6732.

Xing, H., Mo, Y., Liao, W., and Zhang, M.Q. (2012). Genome-wide localization of protein-DNA binding and histone modification by a Bayesian change-point method with ChIP-seq data. PLoS Comput. Biol. 8, e1002613.

Yu, S., Li, J., Ji, G., Ng, Z.L., Siew, J., Lo, W.N., Ye, Y., Chew, Y.Y., Long, Y.C., Zhang, W., et al. (2020). Npac Is a Co-factor of Histone H3K36me3 and Regulates Transcriptional Elongation in Mouse ES Cells. BioRxiv.

Zaidi, S., and Brueckner, M. (2017). Genetics and genomics of congenital heart disease. Circ. Res. 120, 923–940.

Zaidi, S., Choi, M., Wakimoto, H., Ma, L., Jiang, J., Overton, J.D., Romano-Adesman, A., Bjornson, R.D., Breitbart, R.E., Brown, K.K., et al. (2013). De novo mutations in histone-modifying genes in congenital heart disease. Nature 498, 220–223.

Zhang, Y., Zheng, Y., Qin, L., Wang, S., Buchko, G.W., and Garavito, R.M. (2014). Structural characterization of a β-hydroxyacid dehydrogenase from Geobacter sulfurreducens and Geobacter metallireducens with succinic semialdehyde reductase activity. Biochimie 104, 61–69.

Zhu, X., Deng, X., Huang, G., Wang, J., Yang, J., Chen, S., Ma, X., and Wang, B. (2014). A novel mutation of Hyaluronan synthase 2 gene in Chinese children with ventricular septal defect. PLoS One 9, e87437.

