## Supplementary material for "Integration of Protein Interactome Networks with Congenital Heart Disease Variants Reveals Candidate Disease Genes": Methods

### LEAD CONTACT AND MATERIALS AVAILABILITY

### EXPERIMENTAL MODEL AND SUBJECT DETAILS

#### Cell Lines

HEK293T (human [*Homo sapiens*] fetal kidney) and HeLa (human [*Homo sapiens*] cervical cancer) cells were all obtained from ATCC (https://www.atcc.org/). HEK-293 and HeLa cells were cultured in Dulbecco's Modified Eagle Medium (DMEM), high glucose, GlutaMAX™ Supplement (Cat.10566016, Thermo Fisher Scientific) supplemented with 10% fetal bovine serum, 2mM sodium pyruvate, 2mM Non-essential amino acids and 100 I.U./mL penicillin and 100 µg/mL streptomycin.

The WTC11 human iPSC line (Cat. GM25256, Coriell) was obtained from the Gladstone Stem Cell Core (https://labs.gladstone.org/stem_cell). The WTC11 line is used as a normal control by research groups all over the world. This hiPSC line was derived from a skin biopsy from a healthy adult Asian male donor in his early thirties, who showed normal function in a battery of tests. The original fibroblasts were reprogrammed using episomal methods with the following factors: LIN28A, MYC (c-MYC), POU5F1 (OCT4), and SOX2 and pluripotency state and differentiation potential characterized as previously described (Miyaoka et al. 2014).

All hiPSC clones were recurrently verified free of mycoplasma contamination and checked for normal karyotype. Karyotyping analyses were performed by Cell Line Genetics.

#### Human iPS cell line generation by CRISPR-Cas9 editing

To generate the GATA4 and TBX5 knockout lines, WTC11 hiPSCs were dissociated in accutase (Cat. 07920, Stem Cell Technologies), and 250.000 cells were aliquoted per condition and nucleofected with Cas9-ribonucleoprotein complexes (Cas9-RNP) following the Primary Cell Nucleofection P3 Kit manufacturer's instructions (V4XP-3960, Lonza). For Cas9-RNP complex preparation 180pmol of each synthetic modified sgRNA (Synthego) to target exon4 of GATA4 (GAGGCCCACUCGGCGGGAGG) or exon6 TBX5 (GCTTACCTTGTGGTTCTGGTAGG) and 20pmol of SpCas9-NLS purified protein (QB3 Macrolab, UCB) were diluted into 20µl of nucleofection buffer prepared as indicated in Primary Cell Nucleofection P3 Kit, mixed and incubated at room temperature for 10 min. After 10 min. of incubation, the aliquoted 250.000 cells were resuspended in 20µL nucleofection buffer containing the corresponding Cas9-RNP complexes, mixed 5-6 times and then transferred to the bottom of a nucleofector well (nucleofector 96 well cassette from the Primary Cell Nucleofection P3 Kit). This was repeated for every Cas9-RNP condition. The nucleofector cassette was placed in the nucleofector instrument (Nucleofector 4D system, Lonza) and cells nucleofector using the preset program DS-138. The nucleofection cassette was brought back to sterile hood and 80µL of Essential 8 medium (E8) (Cat. A1517001, Life Technologies) with 5µM rho kinase (ROCK) inhibitor (Y-27632 2HCl, Cat. S1049, Selleckchem.com) added into each well and incubated for 10 min. at 37°C. During the incubation time, we removed the hESC-qualified LDEV-free matrigel (Cat. 354277, Corning) from 12 well plates (Corning) pre-coated for 1 hr. at 37°C with 0.5ml per well of matrigel and added 2mL of pre-warmed E8 medium plus 5µM ROCK inhibitor (Y-27632 2HCl) in each well. Then pipetted the nucleofected cells to each of the pre-coted wells with media. After ~3-5 days, wells with surviving clones were expanded to isolate gDNA for screening. A genomic fragment spanning the gRNA target sites was amplified using primers FW 5'AGAGATCTCATGCAGGGTCG3' and REV 5'TCATGATGCCTGGCCTTACT3' for GATA4 with Titanium Taq DNA Polymerase (Cat. 639209, Takara Bio) and primers FW 5'GCAGAAACAGTTGCCCAGAA3' and REV 5'CAAGGCGAATTTAGAGGGCG3' for TBX5 with Phusion® High-Fidelity PCR Master Mix with

GC Buffer (Cat. M0532S, New England BioLabs) and Sanger sequenced (Quintara Biosciences or MCLAB) to identify clones with frame-shift insertions/deletions. Synthego ICE analysis was run to identify clones with highest knock-out efficiency (https://ice.synthego.com/#/), which were subsequently subjected to the colony picking and clone sequencing until monoclonal lines were generated. The top 5 sgRNA predicted off-targets (https://horizondiscovery.com/en/products/tools/crispr-specificity-analysis) were verified to be intact by sanger sequencing in the final monoclonal lines and checked for normal karyotype (Cell Line Genetics).

**Human iPSC culture**

Human iPSCs were maintained on tissue culture-treated polystyrene plates (Cat. 430630, Corning) with hESC-qualified LDEV-free matrigel (Cat. 354277, Corning) in Essential 8 medium (E8) (Cat. A1517001, Life Technologies). The medium was changed daily and the hiPSCs were split every 4 – 6 days using Accutase (Cat. 07920, Stem Cell Technologies). The rho kinase (ROCK) inhibitor (Y-27632 2HCl, Cat. S1049, Selleckchem.com) was included in the medium at 5µM final concentration on the day of passaging.

**Induced cardiomyocyte differentiation**

For human cardiac differentiation into CPs and CMs, we modified the protocols originally developed by Lian et al. and Tohyama et al. to achieve stage-specific, high yield, high-purity cardiac commitment in vitro (Lian et al. 2013; Tohyama et al. 2013). Briefly hiPSCs were detached from hESC-qualified LDEV-free matrigel (Cat. 354277, Corning) with accutase (Cat. 07920, Stem Cell Technologies) and reseeded on matrigel at 0.6–1.2x$10^5$ cells per 12well in Essential 8 medium (E8) (Cat. A1517001, Life Technologies) with 5µM ROCK inhibitor (Y-27632 2HCl, Cat. S1049, Selleckchem.com) (day-3). We optimized each hiPSC clone individually to identify the best cell seeding density that resulted in high levels of cTNT, NKX2-5, TBX5 and GATA4-positive sheet-like beating CMs at day15 of differentiation. On the next two days, media was changed daily with fresh E8 medium without ROCK inhibitor. On the day of cardiac induction (day0), 6µM CHIR99021 (Cat. 4423, Tocris) was added for 24 hr. in 1ml per well of B27-supplemented (without insulin) RPMI1640 media (Cat. 11875-119, Life Technologies). On day1 media was changed for freshly prepared 6µM CHIR99021 in B27-supplemented (without insulin) (Cat. A1895601, Life Technologies) RPMI1640 media. At day 2 and 3 IWP4 (Cat. 5214, Tocris) in B27-supplemented (without insulin) RPMI1640 media was added to activate Wnt signaling at a final concentration of 5µM. At day 4, IWP4 was removed, and 1ml of B27-supplemented (without insulin) RPMI1640 media per was daily added per well (days 4-9). At day 10 the media was changed to regular B27-supplemented (with insulin) (Cat. A1895601, Life Technologies) RPMI1640 hereafter. Typically, in parallel differentiations under identical conditions, WT cells started spontaneous contraction as early as day8 while TBX5-KO and GATA4-KO lines tend to be slightly delayed by 48-96 hr. To ensure consistency across the differentiations used for experimental proposes, from each differentiation cells were collected at day6 and day15 for cTNT, TBX5, NKX2-5 and GATA4 FACS analysis. WT hiPSC-CP samples used for AP-MS corresponded to differentiations with sheet-like beating cardiomyocytes and a minimum of 80% $cTNT^+$, 70% $TBX5^+$, 70% $GATA4^+$, 70% $NKX2\text{-}5^+$ cells analyzed by FACS at day15 of differentiation. Similarly, GATA4-KO and TBX5-KO CP samples corresponded to differentiations with sheet-like beating cardiomyocytes, were TBX5 or GATA4 protein absence had been confirmed, and had a minimum of 40% cTNT positive cells at day15 of differentiation.

**Adenovirus**

Adenoviral - Human Type 5 (dE1/E3) viral particles expressing GLYR1 wild-type (Ad-GFP-EF1-h-GLYR1; PFU titer: $1x10^{10}$ PFU/ml), GLYR1 P496L mutant (Ad-GFP-EF1-h-GLYR1 P496L PFU titer: $2x10^9$ PFU/ml) or negative control (Ad-EF1a-eGFP; PFU titer: $1.2x10^{10}$ PFU/ml) viral particles, were obtained from Vector Biolabs.

### METHOD DETAILS

#### Immunocytochemistry

Media was removed from day 15 CMs and fixed on 12 well plates by adding 1ml 4% formaldehyde (Cat. 28906, ThermoScientific) followed by a 15 min. incubation at room temperature (RT). After the incubation, fixed cells were washed 3 times with 1ml of PBS (without Ca2+ and Mg2+) and stored at 4°C in 1ml of PBS until processed for immunostaining. At first, cells were permeabilized for 45 min. at RT in 1ml of permeabilization/blocking buffer (5% donkey serum, 0.2% triton-X, PBS 1x). CMs were stained overnight at 4°C on gentle agitation with the following indicated primary antibodies: mouse monoclonal anti-Troponin T, Cardiac Isoform Ab-1 Clone 13-11 (REF MS-295-P, Thermo Scientific) (1:100) together with goat polyclonal C-20 anti-TBX5 (Cat. sc-17866, Santa Cruz) (1:50) or goat polyclonal C-20 anti-GATA4 (Cat. sc-1237, Santa Cruz) (1:50) diluted in permeabilization/blocking buffer. The next day wells were washed 3 times with 1ml of PBS per well followed by a 10 min. incubation on gentle agitation at RT. Cells were stained with Donkey Anti-Mouse Alexa Fluor 647 (Cat. A-31571, Thermo Fisher Scientific) and Donkey Anti-Goat Alexa Fluor 488 (Cat. A11055, Thermo Fisher Scientific) secondary fluorophore conjugated antibodies (1:300) for 45 min. on gentle agitation at RT, protected from light exposure. After the incubation, cells were washed 3 times with 1ml of PBS per well followed by a 10 min. incubation on gentle agitation at RT. Immunostained samples were counterstained with DAPI (Cat. 422801, BioLegend) and visualized in a Zeiss Z1 microscope and associated ZEN software.

#### Flow Cytometry

At CP (day6) or CM (day15) stages of differentiation cells were dissociated in accutase (Cat. 07920, Stem Cell Technologies), and fixed in 1.5ml Eppendorf tubes with 1ml of 4% formaldehyde (Cat. 28906, ThermoScientific) for 15 min. on rotation at RT. After the incubation, fixed cells were washed 3 times with 1ml of PBS (without Ca2+ and Mg2+) and stored at 4°C in 1ml of PBS until processed for staining. For staining, cells were pelleted and PBS removed. Cell pellets were resuspended in 200µl of FACS buffer [1x PBS, 5% BSA, 5mM EDTA, 0.25% triton-X] and incubated at RT for 1 hr. The permeabilized cells were then pelleted resuspended in 3µl of staining-A in FACS buffer: mouse monoclonal anti-Cardiac Troponin T [1C11] (1:100) (Cat. ab8295, Abcam) and goat polyclonal anti-NKX2-5 (1:50) (Cat. sc-8697, Santa Cruz); or staining-B: mouse monoclonal A-6 anti-TBX5 (1:50) (Cat. sc-515536, Santa Cruz) and goat polyclonal C-20 anti-GATA4 (1:50) (Cat. sc-1237, Santa Cruz) and incubated at RT for 1 hr. Cells were then washed/spined down 3 times with 200µl of FACS buffer, resuspended in 30µl of FACS buffer with the secondary antibodies diluted 1:300: Donkey Anti-Mouse Alexa Fluor 647 or 568 (Cat. A-31571 or A10037, Thermo Fisher Scientific) and Donkey Anti-Goat 488 Alexa Fluor (Cat. A11055, Thermo Fisher Scientific) and incubated for 1 hr. protected from the light at RT. The stained cells were washed/spined down 3 times with 200µl of FACS buffer and finally resuspended in 200µl of FACS buffer until analyzed. FACS stained samples were measured using FACSCalibur (BD Biosciences) or LSRII (BD Biosciences) and further analyzed using Flowjo software (https://www.flowjo.com/, FlowJo LLC).

**Cardiomyocyte beating characterization**

WT, TBX5-KO and GATA4-KO CMs contractility beat rate parameter was measured from brightfield acquired videos with the Pulse automated analysis software (https://www.pulsevideoanalysis.com). The onset of beating was determined by careful visual daily examination of multiple independent differentiations.

**Plasmid generation: cloning and mutagenesis**

The GFP-GATA4 plasmid (pEN563-pCAGG-eGFP-GATA4) was generated by PCR amplification of the hGATA4 ORF from the precursor vector pAAV2.1CAG-hGATA4 (FW primer: TGGTGGATCCACCGGTATGTATCAGAGCTTGGCCATGG; REV primer:

TGAGCGGCCGCGTTTAAACTTACGCAGTGATTATGTCCCCGTG) and cloned following the Cold Fusion Cloning kit (Cat. MC101B-1, Systems Biosciences). Followed manufacturer's recommendations to linearize the pEN563-CAGG-eGFP vector with PmeI and AgeI (Cat. R0560S and R0552S, New England BioLabs) enzymes to replace the previous contained ORF for the hGATA4. The hGLYR1-MYC and hSMARCC1-MYC plasmids (pCMV-T7-cDNA-MYC-IRES2-mCherry-pA; Cat. EX-Z0806-M73 and EX-A6386-M73) were obtained from GeneCopoeia™. The hGLYR1-MYC P496L was generated with the QuikChange II XL Site Directed Mutagenesis kit (Cat. 200521-5, Agilent Technologies) of the hGLYR1-MYC plasmid with the (FW: GCAAGGAAACTTTAAGCTAGATTTCTACCTGAAATAC and RV: GTATTTCAGGTAGAAATCTAGCTTAAAGTTTCCTTGC). The HA-hBRD4 plasmid (p6344 pcDNA4-TO-HA-Brd4FL) was a gift from Peter Howley (Addgene plasmid # 31351; http://n2t.net/addgene:31351; RRID: Addgene_31351) (Rahman et al. 2011).

**Nuclear enriched lysis, Immunoprecipitation and Immunoblotting**

WT, GATA4-KO and TBX5-KO hIPSC derived CPs or HEK293 cells transfected with the indicated expression vectors following manufacturer instructions (FuGene HD, Promega) were lysed in Cell Lysis buffer [20mM Tris-HCl pH 8, 85mM KCl, 0.5% NP-40, freshly added protease and phosphatase inhibitors (Cat. 4693132001 Roche and Cat. 4906837001 Sigma-Aldrich)] and incubated on rotator at 4°C for 10 min. Cells were spun down at 2500 rpm for 5min. at 4°C to pellet nuclei and supernatants containing the cytosolic fraction were removed and stored for quality control proposes. Nuclei were resuspended in Nuclear Extraction Buffer (NEB) [20 mM HEPES, pH 7.4, 0.5 M NaCl, 2 mM MgCl2, 1 mM CaCl2, 0.5 % NP-40, K-Acetate 110mM, 1µM ZnCl2, and freshly added Benzonase 2µl enzyme/ml buffer (Cat. E1014, Millipore), protease and phosphatase inhibitors] and incubated on rotation for 30 min. at 4°C. For CPs, 600µl NEB were used per 4x 12 well plates of initial CPs (~$100x10^6$), whereas the nuclei resulting from one 10 mm confluent HEK293 plate were lysed in 300µl of NEB. After incubation, samples were centrifuged at max speed for 10 min. and the nuclear enriched lysates (supernatants) were moved to clean tubes. Nuclear lysates were then diluted 1:3 in Nuclear Dilution Buffer (NDB) [20 mM HEPES, pH 7.9, 1 mM EDTA, 0.2 % NP-40, freshly added protease and phosphatase inhibitors], 1200µl of NDB for CPs; 600µl NDB for HEK293. At this step samples can be stored and later used for immunoprecipitation or for western blotting.

The immunoprecipitations (IP) were done as described before (González-Terán et al. 2016) with modifications. Briefly, 50µl of Dynabeads™ Protein G magnetic (Cat. 10004D, Invitrogen) per IP sample were washed twice with 1ml PBS using a magnetic stand, and resuspended in an equivalent volume to the original volume of beads of PBS. Magnetic beads were then conjugated with a primary antibody: per 50µl of magnetic Dynabeads 4µg of mouse anti- GATA-4 Antibody (G-4) (Cat. sc-25310 X, Santa Cruz), 4µg mouse anti- TBX5 Antibody (A-6) (Cat. sc-515536 X, Santa Cruz), 2µg of anti-Myc tag antibody - ChIP Grade (Cat. ab9132, Abcam) or 2µl of GLYR1 anti-sera 7 (James T. Kadonaga Laboratory) were added and incubated on rotation for 1 hr. at 4°C. Then, the extra non-conjugated antibody was removed by washing the incubated Dynabeads with 1ml of PBS and twice with 1ml of Nuclear Dilution Buffer (NDB) using the magnetic stand, and resuspended in the initial volume of NDB. At this point the coated Dynabeads are ready for IP. Protein content was quantified by Quick Start™ Bradford Protein Assay Kit 1 (Cat. 5000201, BioRad) from nuclear enriched lysates and 1mg (immunoblot) or 3mg (mass spectrometry) of total protein per IP of endogenous proteins or 150µg (immunoblot) or 1mg (mass spectrometry) per IP of ectopically expressed proteins were aliquoted per condition and volumes were equalized with NDB. Prior to adding 50µl of coated beads per IP sample, 50µl of the prepared nuclear enriched lysates were set aside as “inputs”. IP samples were incubated with coated Dynabeads 4 hrs. for endogenous IP or 1 hr. for IP of overexpressed proteins, on rotator at 4°C. After incubation, samples were placed in the magnetic stand and supernatants removed and saved as “unbound-fraction” for quality control proposes. Beads were washed 2x with 1ml of NDB, and 3 times with 1ml of NDB buffer without detergent. After the last wash, beads were spun down and tubes put on the magnet to remove the remaining liquid. At this point the IP samples can be processed for mass spectrometry or subjected to immunoblotting.

For immunoblotting, samples were subjected to PAGE-SDS. Firstly, 1x of NuPAGE™ LDS Sample Buffer (4X) (Cat. NP0007, Thermo Fisher Scientific) was added to IP samples or to 25-50µg of nuclear enriched lysates and boiled at 95°C for 10 min. and 5 min. respectively. Samples were resolved in pre-cast Novex 4-12% Tris-Glycine gels (Cat. XP04122BOX, Invitrogen) and transferred over night at 35V into polyvinylidene difluoride (PVDF) membranes for endogenous proteins. For ectopically expressed proteins gels were transferred using iBlot® Transfer Stack, PVDF, mini (Cat. IB4010-32, ThermoScientific) and iBlot™ gel transfer device (Thermo Fisher Scientific) for 8 min. at 200V. Membranes were blocked with LI-COR blocking buffer (Cat. 927-40010, LI-COR Biosciences) or 10% BSA in PBS-T (PBS with 0.1% Tween) for 20 min., and incubated overnight on agitation with the primary antibody. Appropriate secondary HRP-conjugated antibody (Abcam) or fluorophore conjugated secondary antibody (LI-COR) was added for 1 hr. at a dilution of 1:5000 followed by detection with ECL Prime Western Blotting Detection Reagent (Cat. RPN2232, GELife Sciences) and exposure to autoradiography film at various time intervals or by digital imaging (LI-COR Odyssey). The following primary antibodies were used for immunoblotting: mouse anti- GATA-4 Antibody (G-4) (Cat. sc-25310 X, Santa Cruz), mouse anti-TBX5 Antibody (A-6) (Cat. sc-515536 X, Santa Cruz), anti-Myc tag antibody - ChIP Grade (Cat. ab9132, Abcam), GLYR1 anti-sera 7 (James T. Kadonaga Laboratory), anti-HA tag antibody - ChIP Grade (Cat. ab9110, Abcam) and anti-Vinculin Monoclonal Antibody VLN01 (Cat. MA5-11690, Thermo Fisher Scientific).

**Mass Spectrometry**

For mass spectrometry all the IP steps described were processed in protein low-bind sterile Eppendorf tubes (Cat. 022431102, Eppendorf), aliquots specific for mass spectrometry were made for each of the IP buffers and carefully managed to avoid contaminations with filter tips.

After the magnetic beads were spun down and the excess of liquid removed, to each IP sample tube another 1ml of NDB was added, beads resuspended and transferred using wide orifice tips to new protein low-bind sterile Eppendorf tubes. IP samples were returned to the magnetic stand and supernatant carefully removed by aspiration. IP samples were spun down again to collect the remaining liquid, placed in the magnet and the remaining liquid removed. Next, we proceeded with on-bead protein digestion. One bead volume (25µl) of freshly prepared Alkylation Buffer [2M Urea; 50mM Tris, pH8.0; 1mM DTT; 3mM IODO; resuspended in LC-MS high grade water] was added to each IP sample and incubated at RT in the dark for 45 min. while shaking to ensure bead suspension. After incubation, an additional 3mM DTT and 750ng of trypsin per 10µl of bead volume were added to each tube. IP samples were on-bead digested overnight at 37°C on agitation. The next morning, beads were pelleted in a microcentrifuge at 2000 rpm for 4 min. IP samples were placed in a magnetic stand and supernatants carefully transferred to a fresh 0.5ml protein lo-bind tubes. Beads can be saved at 4°C for quality control. To each 0.5ml tube containing the digested IP supernatants, formic acid at a final concentration of 1% was added to stop the digestion process. The processed samples can be stored at this point at -80°C until desalting. Samples were subjected to desalting with OMICS tips (Agilent) and lyophilization in a speed-vacuum concentrator for 30 min. Lyophilized samples were stored at -20°C until proceeding with mass spec analysis. When ready, lyophilized samples were resuspended in 10µl of 0.2% formic acid/ 2%acetonitrile immediately before loading into the mass spectrometry instrument. All APMS samples were measured using the Q Exactive Hybrid Quadrupole-Orbitrap™ Mass Spectrometrer.

**RNA extraction, RT-PCR and real-time PCR analysis**

Cells were harvested in TRIzolTM LS reagent (Cat. 10296010, Invitogen) and total RNA was extracted using the Direct-Zol RNA kit (Cat. R2052, Zymo Research) according to manufacturer instruction. 1000ng of RNA were converted to cDNA using SuperScriptTM III First-strand

Synthesis SuperMix for qRT-PCR (Cat. 18080400, Invitrogen). For Taqman real-time PCR, 1/50 cDNA was applied for quantitative PCR reaction using Taqman Universal PCR master mix (Cat. 4305719, Life technologies). The PCR was conducted in 7900HT Fast Real-Time system (Applied Biosystem). The Taqman probe for enhanced GFP (eGFP) quantification (Mr04097229_mr, ThermoFisher Scientific). All gene expressions were normalized with human GAPDH levels (Hs99999905_m1, ThermoFisher Scientific).

**RNAseq Assay**

Total RNA was TRIzol-extracted (Cat. 10296010, Thermo Fisher Scientific) and further purified using the Direct-Zol RNA kit (Cat. R2052, Zymo Research) with DNaseI in-column treatment according to the manufacturer's instructions and quantified with Nanodrop (Thermo scientific). After RNA quality control with bioanalyzer Agilent 2100 (Agilent Technologies), Paired-end Poly(A)-enriched RNA libraries were prepared with the Ovation RNA-seq System V2 Kit (Cat. 7102-08, NuGEN; strand specific) from the Gladstone Genomic core. The mRNA-seq libraries were analyzed by Agilent Bioanalyzer and quantified using an Illumina Library Quantification Kit (Cat. KK4824, KAPA Biosystems). Libraries were prepared by the Gladstone Genomics Core (http://labs.gladstone.org/genomics/home). High-throughput sequencing was done using an Illumina HiSeq 2500 instrument (http://humangenetics.ucsf.edu/genomics-services/sample-processing/).

**ChIPseq assay, library preparation and sequencing**

For ChIPseq experiments, CPs ($30\times10^6$ for cTFs and $10\times10^6$ for GLYR1 and histone marks) were pelleted and suspended in 10ml DMEM and cross-linked in 1% Formaldehyde solution (Cat. 28906, Thermo Fisher Scientific) by rocking in room temperature for 10 min. Then glycine (final concentration 0.125M) was added to quench the cross-link for 5 min. Samples were centrifuged at 1000 rcf for 5 min. at 4°C. Cells were washed with 10ml of cold 1x PBS supplemented with proteinase inhibitors and phosphatase inhibitors (Cat. 4693132001 Roche and Cat. 4906837001 Sigma-Aldrich) and the pellets were snap frozen in liquid nitrogen. All samples were stored at −80°C until use. When ready, cell pellets were incubated in cell lysis buffer (20 mM Tris-HCl, pH 8, 85 mM KCl, 0.5% NP-40, protease/phosphatase inhibitors) for 10 min. on a rotator at 4°C. Nuclei were isolated by centrifugation (2,500 x g, 5 min., 4°C), resuspended in nuclear lysis buffer (50 mM Tris-HCl, pH 8, 10 mM EDTA, pH 8, 1% SDS, protease/phosphatase inhibitors) and incubated on a rotator for 30 min. at 4°C. Chromatin was sheared using a Covaris S2 sonicator (Covaris Inc) for 15 min. (60 s cycles, 5% duty cycle, 200 cycles/burst, intensity = 6) until DNA was in the 200–700 base pair range. Chromatin was diluted 3-fold in ChIP dilution buffer (0.01% SDS, 1.1% Triton X-100, 1.2mMEDTA, 16.7mMTris-HCl, pH 8, 167 mM NaCl, protease/phosphatase inhibitors) and incubated with the corresponding primary antibody at 4°C overnight under rotation. Antibody-protein complexes were immunoprecipitated using 50μl of Dynabeads™ Protein A/Protein G (Cat. 10015D, Invitrogen) per sample at 4°C for 2 h under rotation. After incubation, beads were washed five times (2 min./wash under rotation) with cold RIPA buffer [50 mM HEPES-KOH, pH 7.5, 500 mM LiCl, 1 mM EDTA, 1% NP-40, 0.7% Na-deoxycholate], followed by one wash in cold final wash buffer [1xTE, 50 mM NaCl]. Immunoprecipitated chromatin was eluted at 65°C with agitation for 30 min. in elution buffer [50mMTris-HCl pH 8.0, 10mMEDTA, 1% SDS]. High-salt buffer [250mM Tris-HCl, pH 7.5, 32.5 mM EDTA, pH 8, 1.25M NaCl] and Proteinase K (Cat. P8107s, New England Biolabs Inc (NEB)) were added and crosslinks were reversed overnight at 65°C. Samples were treated with RNase A, and DNA was purified with AMPure XP beads (Cat. A63881, Beckman Coulter). Fragmented ChIP and input DNA were end-repaired, 5'-phosphorylated and dA-tailed with NEBNext Ultra II DNA Library Prep Kit for Illumina (Cat. E7645, New England BioLabs). Samples were ligated to adaptor oligos for multiplex sequencing (Cat. E7335, New England BioLabs), PCR amplified, and sequenced on an Illumina NextSeq 500 at the Gladstone Institutes. Primary antibodies used for ChIP were: GATA4 (Cat. sc-1237 X, Santa Cruz), TBX5 (Cat. sc-17866 X, Santa Cruz),

H3K36me3 (Cat. ab9050, Abcam), GLYR1 (anti-serum 7; provided by James T. Kadonaga's laboratory), NKX2-5 (Cat. sc-8697 X, Santa Cruz), MEIS1 (Cat. ab19867, Abcam), ISL1 (Cat. AF1837, R&D Systems). The specificity of the antibodies was validated in previous publications (Luna-Zurita et al. 2016; Dupays et al. 2015; Fei et al. 2018).

**siRNA transfection on CPs**

For siRNA knockdown experiments during in cardiac progenitor cells, at day 4 of differentiation cells were detached from the plates with 1ml of accutase (Cat. 07920, Stem Cell Technologies) and quenched with 1ml of B27 (minus insulin) RPMI1640 media (Cat. 11875-119, Life Technologies) per well. All cells were combined and centrifuge at 300xg, supernatant removed and the pellet resuspended in a volume of B27 (minus insulin) RPMI with 5uM ROCK inhibitor (Y-27632 2HCl, Cat. S1049, Selleckchem.com) necessary for 2x the number of wells initially collected. Cells were then seeded in twice the number of 12 well plates originally collected that were pre-coated with fibronectin bovine plasma solution (Cat. F1141, Sigma-Aldrich). Cells were immediately transfected in solution, prior attachment to the well surface using lipofectamine RNAiMax (Cat. 13778075, Invitrogen). For one well of a 12 well plate, mix A (75µl Opti-MEM (Cat. 31985070, Thermo Fisher Scientific) were combined with 3µl of a 10µM siRNA stock) and mix B (75µl Opti-MEM) with 7µl of lipofectamine RNAiMax were prepared. Mix A and B were combined and incubated at RT for 5-10 min. 160µl of lipofectamine siRNA complexes were added dropwise to each well. At day 7 of differentiation, ~72 hrs. after transfection cell were collected, washed, supernatants removed and pellets snap frozen and stored at -80°C until processed. The following siRNA were used: GATA4 Silencer Select Pre-designed SiRNA (Cat. 4392420, ID s535120, Lot. AS02F2E2, Thermo Fisher Scientific), siGLYR1 (siRNA ID: SASI_Hs01_00116796, Millipore-Sigma) and Silencer Select Negative control #1 siRNA (Cat. 4390843, Thermo Fisher Scientific).

**Luciferase assay**

GATA4–GLYR1 transcriptional synergy reporter assay was performed using the pANF638L vector (Knowlton et al. 1991) or pGL4.23[luc2/minP] (Cat. E8411, Promega) modified reporters in which putative intronic RE co-bound by GATA4 & GLYR1 were cloned. Briefly, HeLa cells were cultured in 24-well plates at 10^5 cells per well and transfected within 24 hrs. of seeding. Cells were co-transfected with 200 ng of luciferase reporter vector and 20 ng of Renilla luciferase control vector pRL-TK (Cat. E2241, Promega) in 2.4 µl FuGENE HD (Cat. E2311, Promega) and 43 µl Opti-MEM (Cat. 31985070, Thermo Fisher Scientific). The transfection mix was aliquoted in 5 tubes (1 per condition) and the following conditions were prepared for the luciferase assay using the pANF638L vector: 1.) Control: 600ng of empty vector (EV); 2.) GATA4: 200ng of GFP-GATA4 vector plus 400ng EV; 3.) GLYR1: 400ng MYC-GLYR1 vector plus 200ng EV and 4.) GATA4+GLYR1: 200ng of GFP-GATA4 vector with 400ng MYC-GLYR1 vector and 5.) GATA4+GLYR1 P496L: 200ng of GFP-GATA4 vector with 400ng MYC-GLYR1 P496L vector. Media was changed 24 hrs. after transfection and cells collected 48 hrs. following transfection. Samples were processed with the Dual Luciferase Assay System (Cat. E1960, Promega) following manufacturer's instructions and measured with a luminometer (SpectraMax i3). The GATA4–BRD4 transcriptional synergy was tested in three technical replicates in 3 independent experiments.

For analyzing the GATA4–GLYR1 transcriptional synergy within putative intronic REs, GATA4 -bound intronic regions within GATA4 & GLYR1 -bound cardiac development genes that co-localized with H3K27ac, H3K4me1 or H3K4me3, MED1 and at least co-occupied by two cTFs, were cloned with Cold Phusion (Systems Biosciences) by designing gBlocks (IDT) for each of the selected putative REs (GATA6: chr18:19,773,958-19,774,419; MYL4: chr17:45,296,131-45,296,972; TTN: chr2:179,493,152-179,494,179 ) flanked by homology arms complementary to the pGL4.23[luc2/minP] luciferase reporter vector (Cat. E8411, Promega). HeLa cells were plated as indicated for the pANF638L reporter and transfection mixed prepared as indicated above. The transfection mix was aliquoted in 5 tubes (1 per condition) and the following conditions were prepared for each of the cloned pGL4.23 luciferase reporter vectors: 1.) Control: 200ng of empty vector (EV); 2.) GATA4: 200ng of GFP-GATA4 vector plus Adenovirus control (Ad-EF1a-eGFP, MOI 25); 3.) GLYR1: Adenovirus GLYR1 WT (Ad-eGFP-EF1-h-GLYR1, MOI 25) and 4.) GATA4+GLYR1: 200ng of GFP-GATA4 vector and Adenovirus GLYR1 WT (MOI 25); 5.) GATA4+GLYR1 P496L: 200ng of GFP-GATA4 vector Adenovirus GLYR1 P496L (Ad-eGFP-EF1-h-GLYR1 P496L, MOI 15). The MOIs were determined by HeLa cell infection followed by quantitative PCR amplification with the eGFP Taqman probe (Mr04097229_mr, ThermoFisher Scientific). MOIs rendering comparable eGFP expression levels were chosen for the luciferase experiments (Relative to GAPDH Avg levels for Ad GLYR1 WT MOI 25: 50.134 and Ad GLYR1 P496L MOI 15: 57.333; n=3). HeLa cells were collected 48h after transfection/infection and processed as indicated for the pANF638L vector.

### COMPUTATIONAL ANALYSIS

#### Mass spectrometry analysis of affinity purifications.

Peptides from affinity purifications were analyzed on a Q-Exactive Plus (Thermo Fisher) mass spectrometer. The Q-Exactive Plus system was equipped with an Easy1200 nLC system (Thermo Fisher) and an analytical column (25 cm x 75 um I.D. packed with ReproSil Pur C18 1.9 µm, 120Å particles, Dr. Maisch). A gradient was delivered from 2% to 30% acetonitrile over 53 minutes at a flow rate of 300 nl/min. All MS spectra were collected with Orbitrap detection, while the 20 most abundant ions were fragmented by HCD and detected in the Orbitrap. Peptide and protein identification searches, as well as label-free quantitation were performed using the MaxQuant data analysis algorithm (version 1.5.8.0) (Cox and Mann 2008). Data were searched against a database containing SwissProt Human sequences (downloaded 02/2017) concatenated to a decoy database where each sequence was randomized in order to estimate the false discovery rate (FDR).

Variable modifications were allowed for methionine oxidation and protein N-terminus acetylation. A fixed modification was indicated for cysteine carbamidomethylation. Full trypsin specificity was required. The first search was performed with a mass accuracy of +/- 20 parts per million (ppm) and the main search was performed with a mass accuracy of +/- 4.5 parts per million. A maximum of 5 modifications were allowed per peptide. A maximum of 2 missed cleavages were allowed. The maximum charge allowed was 7+. Individual peptide mass tolerances were allowed. For MS/MS matching, a mass tolerance of +/- 20 ppm was allowed and the top 12 peaks per 100 Da were analyzed. MS/MS matching was allowed for higher charge states, water and ammonia loss events. The data were filtered to obtain a peptide, protein, and site-level false discovery rate of 0.01. The minimum peptide length was 7 amino acids.

**Selection of Interactome Proteins**

APMS data was analyzed using the artMS package (D. Jimenez-Morales et al. 2020) in R followed by protein-protein interaction scoring by the SAINTq software (Teo et al. 2016) to identify significantly-interacting proteins for GATA4 and TBX5 baits. Default parameters for both softwares was used except where indicated here: To create the GATA4 interactome, we analyze at the protein level and select proteins that interact at a BFDR cutoff of <= 0.001; to create the TBX5 interactome, we analyze at the peptide level and select those that interact at a BFDR cutoff of <= 0.05. Intensity data from the control (knockout) cell lines was normalized per SAINTq configuration options such that the average total intensity in each bait purification was equal to the average total intensity across the control experiments.

To focus on transcriptionally-relevant interactions, we additionally filter proteins by those that appear in the nuclear compartment, those that are expressed at detectable levels in at least one of the same cell types as the bait, and proteins whose gene expression was significantly lower in the control line but did not have a greater than 0.5 log-fold change drop in intensity. Nuclear compartment genes were identified using the Cytoscape package BiNGO (Maere et al. 2005; Shannon et al. 2003) with additional manual curation from literature. For cell type expression, single-cell RNA-seq data from deSoysa et al. 2019 was used to determine if an interaction was likely to occur, given co-expression in the same cell type. Briefly, mesoderm and neural crest cells in the developing heart were used to identify seven cell type populations (multipotent Isl1+ progenitors, endothelial or endocardial cells, epicardium, myocardium, neural crest-derived mesenchyme, paraxial mesoderm and lateral plate mesoderm) (de Soysa et al. 2019); proteins were considered to be potentially physiologically relevant interactors if they were detected at any level in one of the same cell types as the bait. Finally, protein hits that were considered more likely to be false positives based on lower expression in the control cell lines, without concomitant reduction in protein intensity, were removed from the interactome list. Significant differential gene expression was determined in R using the edgeR package (Robinson et al. 2010); normalized protein intensities were averaged in all control experiments and bait experiments, and proteins with significantly reduced expression in control with less than a 0.5 log-fold change drop in intensity were not considered to be interactors.

**Gene expression tissue distribution and specificity**

The categories for gene expression tissue distribution and tissue specificity defined by the *Tissue Atlas* within the *Human Protein Atlas* were used to classify the specified gene groups (https://www.proteinatlas.org/humanproteome/tissue/tissue+specific). These classifications are based on transcriptomics analysis across all major organs and tissue types in the human body, where all putative 19670 protein coding genes have been classified with regard to abundance and distribution of transcribed mRNA molecules (Uhlén et al. 2015).

Specificity illustrates the number of genes with elevated or non-elevated expression. Elevated expression includes three subcategory types:

- Tissue enriched: At least four-fold higher mRNA level in a particular tissue compared to any other tissues.
- Group enriched: At least four-fold higher average mRNA level in a group of 2-5 tissues compared to any other tissue.
- Tissue enhanced: At least four-fold higher mRNA level in a particular tissue compared to the average level in all other tissues.

Distribution, on the other hand, visualizes how many genes that have, or do not have, detectable levels (NX≥1) of transcribed mRNA molecules. All elevated genes are categorized as:

- Detected in single: Detected in a single tissue
- Detected in some: Detected in more than one but less than one third of tissues
- Detected in many: Detected in at least a third but not all tissues
- Detected in all: Detected in all tissues

**Permutation-based test**

We tested the adjusted odds ratio of observing a de novo mutation in an interactome gene in CHD probands relative to controls, using data published in Jin et al. 2017 (Jin et al. 2017). We ran 10,000 permutations in which case/control status was randomly shuffled to generate a null distribution of odds ratios. This was performed for non-synonymous de novo mutations, synonymous de novo mutations, and rare inherited loss-of-function mutations (at minor allele frequency $10^{-5}$) (Jin et al. 2017) on the GATA4 and TBX5 interactomes generated from both cardiac progenitor and HEK293T APMS experiments. We observed that some genes appeared to have been more deeply sequenced in control individuals, while other genes showed the opposite trend. This is not unexpected, as control individuals in the Jin et al. dataset were sequenced for a different study and at different institutions from PCGC individuals. Therefore, to control for regional biases in sequencing between the case and control studies, we adjusted the odds ratios of the synonymous and non-synonymous data by a factor that restricts the synonymous odds ratio to 1 (the null expectation). This correction was performed for the observed odds ratio and the odds ratios calculated in each permutation of *de novo* variants. To determine whether this signal was driven by already-identified CHD risk genes, we repeated the analysis after removing de novo variants occurring in known genes (sourced from Jin et al. 2017, Supplementary Data Set 2: 253 Curated known Human/Mouse CHD genes, reproduced in this publication in Table S4) (Jin et al. 2017).

**Diagnostic model**

Fyler Diagnoses of the PCGC probands were sorted into 15 possible categories, chiefly distinguished by developmental stage and/or cell type at which cardiogenesis is disrupted (Table S6). For each proband diagnosed with one of the Fyler diagnoses in a category, we recorded the number of de novo mutations occurring in interactome genes. We used a poisson regression model (implemented using the glm() function in R) to determine whether there was a significant relationship between a given category and the number of interactome genes harboring de novo mutations in each proband in that category. It should be noted that many probands had complex heart defects that spanned multiple diagnoses and categories, and therefore these categories are not mutually exclusive.

**Interactome CHD Candidate Gene characterization**

All de novo variants observed in CHD probands and matched controls were assessed for the following properties: CADD score, pLI score, variant degree, CHD-gene degree, heart expression percentile rank, haploinsufficiency, and number of mutations per kilobase. The residue-level CADD score (Rentzsch et al. 2019) estimates the likely deleteriousness of a variant based on conservation data. pLI score indicates the predicted loss-of-function intolerance of the gene, scaled between 0 and 1, and was sourced from gnomAD version 2.1.1 (Karczewski et al. 2020). Similarly, haploinsufficiency predicts the deleteriousness of having only a single functional copy of a gene. We use the predicted haploinsufficiency values from Huang et al. 2010 (Huang et al. 2010). The CHD-gene degree counts the number of protein-protein interactions that the gene shares with previously-identified CHD risk genes, while the variant degree counts the number of protein-protein interactions shared with other genes that had de novo variants in a CHD proband. These node degree counts were normalized by the total number of connections observed in the gene, and are based on known mammalian protein-protein interactions in iRefIndex version 15.0 (Razick et al. 2008). Finally, the number of mutations per kilobase measures the number of times a de novo or rare loss of function variant was observed in a CHD proband, normalized by the coding length of that gene. We use a Mann-Whitney U test with Bonferroni correction to assess whether interactome genes and genes with non-synonymous de novo mutations differ significantly in these properties.

**Variant scoring**

All non-synonymous de novo variants occurring in GT-interacting genes and observed in CHD probands were ranked based on a series of gene-level, residue-level, and patient-level properties. The observed/expected (o/e) score indicates how often a variant of this type (missense or loss-of-function) was observed in the gene relative to null expectation. Since lower o/e scores indicate higher deleteriousness, this category was ranked descending (such that the highest value received a rank of 1). pLI score indicates the predicted loss-of-function intolerance of the gene, scaled between 0 and 1 where 1 is more intolerant. pLI and o/e data was sourced from gnomAD version 2.1.1 (Karczewski et al. 2020). CHD-gene degree, variant degree, and mutations per kilobase values were calculated as described above (see Methods: Interactome CHD Candidate Gene Characterization). Expression specificity was calculated using data from median transcripts-per-million (tpm) as published in GTEx version 8.1.1.9 (GTEx Consortium et al. 2017).

Average median tpm was calculated for heart tissues (adult atrium, adult left ventricle) and all other available tissues with the exception of testis. The specificity score is then defined as the average tpm in heart tissues normalized by average tpm across all tissues.

For each of these properties, the variants were ranked based on their relative scores. Ties were resolved by taking the average value of the would-be ranks. Missing data was imputed to the median value of the given property. Gene-level rankings (pLI score, CHD-gene degree, variant degree, mutations per kilobase, and expression specificity) and residue-level rankings (oe score and CADD score) were separately averaged and then added together. This average rank sum was then additionally weighted by two factors to capture aspects of their proband-level and protein contexts.

Firstly, if the proband had additional mutations in other interactome genes or other previously-identified CHD genes, we reduced the variant's weight. Specifically, we multiply the rank-sum score by the lowest-applicable factor if they meet any of these conditions:

| Factor | Conditions |
|---|---|
| 0.75 | Proband has another rare (MAF $10^{-5}$) inherited loss-of-function OR missense de novo variant in an interactome gene |
| 0.50 | Proband has a predicted-damaging de novo mutation in an interactome gene or rare inherited loss-of-function mutation in a previously-identified CHD gene |
| 0.25 | Proband has a de novo missense mutation in a previously-identified CHD gene |
| 0.10 | Proband had a de novo missense mutation in a previously-identified CHD gene, and that variant was predicted-damaging or led to protein loss-of-function. |

To summarize, the variant is down-weighted in cases where it is likely that another mutation in the proband is causing or contributing to the CHD phenotype.

Secondly, if the de novo variant leads to protein loss-of-function, or if it occurred in a known protein domain (and therefore is suspected to interfere with protein activity), the variant rank-sum was transmitted as-is. Otherwise, the variant's rank-sum was multiplied by 0.5.

In summary, this framework prioritizes de novo variants that occur in interactors of GATA4 and TBX5, that are more likely to disrupt protein function, and that are more likely to be high-effect monogenic causes of disease.

**GLYR1 Model Organisms Alignment**

Sequences of several vertebrate model organisms containing the rigid loop (bridging the two tetramerization alpha helice bundles) of the NPAC proteins' dehydrogenase domain were aligned using CLC Sequence Viewer 8.0. Amino acids 490-529 were aligned, partially spanning exons 14 and 15 (490-495, 496-5229 respectively) in the *H. sapiens* sequence. Alignments were created with the "Alignment" function with a gap open cost of 10.0, gap extension cost of 1.0, end gap cost as any other, and the very accurate (slow parameter).

Used mRNA (NM) and predicted mRNA (XM) Sequences

Chimp (*Pan Traglodyte*): XM_016929357

Gorilla (*Gorilla gorilla*): XM_019012358

Human (*Homo sapien*): NM_032569

Mouse (*Mus musculus*): NM_001359747.1

Rat (*Rattus norvegicus*): NM_001007800

Chicken (*Gallus gallus*): NM_001006572

Frog (*Xenopus laevis*): NM_001030494

Zebrafish (*Danio rerio*): XM_005164104

**GLYR1 Structural Model**

The wildtype GLYR1 structure was imported from the RCSB Protein Data Bank, entry 2UYY, the structure being elucidated through X-ray diffraction (Tickle, J. et al. 2007). The structure and domains of the NPAC monomer were edited using the PYMOL Molecular Graphics System Version 2.3.5. Domains of the NPAC dehydrogenase domain are defined as by Zhang et al 2014

(Zhang et al. 2014). The protein is shown through the cartoon function, displaying the general tertiary structure of the protein. Amino acid proline 496 and its mutant proline 496 leucine are shown in gold and through the stick function displaying the secondary structure of the amino acids to delineate the significance in change of structure. In the focused images of the rigid loop of the alpha helices tetramerization bundle, the amino acids are again shown through the stick function to delineate the secondary structure interactions of the amino acids.

**Molecular structural dynamics methods**

The initial protein structure for all-atom MD simulations in explicit water of NPAC was downloaded from the Protein Data Bank, code 2uyy.pdb.

Missing atoms and side chains where added using the Protein Preparation Wizard of the Maestro Suite of Programs (v. 2019−4). Proline to Leucine mutation was also performed using Maestro (Maestro Schrödinger, LLC 2019; Sastry et al. 2013). The simulations were run using the same protocol for both the WT and mutated monomer (subunit A).

All systems where allowed to relax with 2000 steps of steepest descent followed by another 2000 steps of conjugate gradient energy minimization. The temperature of the systems was gradually raised to 300 K in the NVT ensemble in 1.2 ns at 1 fs time-step, using the Langevin thermostat. In particular, six runs of 200 ps were performed increasing the temperature of 50 K at each step (T = 50, 100, 150, 200, 250, and 300 K, respectively). At 300 K, the density of the system was adjusted with 1 ns at 2 fs time-step under NPT conditions by weak coupling to a bath of constant pressure ($P_0$ = 1 bar, coupling time $t_p$ = 0.5 ps). The production runs were thus carried out in the NVT ensemble. Bonds involving hydrogen atoms were constrained with the SHAKE algorithm (Miyamoto and Kollman 1992), allowing a time step of 2 fs. Electrostatic forces were computed using the particle mesh Ewald algorithm with a truncation cut-off of 10A° (Darden et al. 1993). The initial velocity of all atoms was obtained from a Maxwellian distribution at the initial temperature of 300 K.

MD simulations where run in 3 independent replicas of 500 ns each (1.5 μs in total per system), Specifically, MD simulations were performed using Amber18 pmemd.CUDA with the all atom ff14SB force field under periodic boundary conditions (Case et al. 2017). The triclinic simulative box, filled with TIP3P (Jorgensen et al. 1983) water molecules and rendered electroneutral by addition of Na+ counterions consists of a final number of atoms of about 41 300 (monomer WT and P496L mutant), particles for each system.

The atomic positions were saved every 10 ps. The equilibrated parts of the trajectories were used for subsequent analyses. Equilibration of the trajectories was checked by monitoring the equilibration of the RMSD with respect to the initial structure and of the internal protein energy. The equilibrated parts of each trajectory for the two systems were next combined into a meta-trajectory, which was subsequently used for all the reported characterizations. Classical structural analyses were carried out with the tools in the Amber18 and Gromacs 4.5.5 package (Bekker et al. 1993) or with code written in-house.

The *root mean square deviation* (RMSD) of the backbone of the protein with respect to first frame of the trajectory along the simulation time has been calculated by least-square fitting the structure to the reference structure ($t_2 = 0$) and subsequently calculating the RMSD

$$\mathrm{RMSD}(t_1,t_2) = \sqrt{\left[\frac{1}{M}\sum_{i=1}^{N} \mathrm{m_i} \parallel \mathrm{r_i}(t_1) - \mathrm{r_i}(t_2) \parallel^2 \right]}$$

where $M = \sum_{i=1}^{N} m_i$ and $r_i(t)$ is the position of atom *i* at time *t*.

The *RMSF* which is a measure of the displacement of each residue averaged over the number of atoms considered, has been calculated relative to the average structure, in the equilibrated part of the simulation.

**Differential gene expression**

In order to identify genes differentially expressed between WT CP versus GATA4-KO or TBX5 CPs (n=5); siControl vs siGATA4 CPs (n=3) and siControl.2 vs siGLYR1 CPs (n=2), the analyses start with raw reads/sequences in FASTQ format. Trimming of known adapters and low-quality regions of reads was performed using Fastq-mcf (Aronesty 2013). Sequence quality control was assessed using the program FastQC (Andrews 2007) and RSeQC (Wang et al. 2012). Alignment of the provided samples to the reference genome was performed using STAR 2.5.2a (Dobin et al. 2013). Reads were aligned to the human hg19 reference assembly indicated in the header of the differential expression file. Reads were assigned to genes using *featureCounts* (Liao et al. 2014), part of the Subread suite (http://subread.sourceforge.net/). Gene-level counts were arrived at using Ensembl gene annotation, in GTF format. Differential expression was assessed using edgeR (Robinson et al. 2010), an R package available through Bioconductor. Genes where there were not at least two samples with at least 5 (raw) reads were filtered out from further analyses. The reads counts of remaining ones are normalized for sample-to-sample variation using *calcNormFactors* in edgeR (Robinson et al. 2010). The mean gene expression was modeled as a function of siRNA status (siRNA treatment vs scramble control) and sample id. Genes whose expression is associated with siRNA status were determined by the likelihood ratio test (Smyth 1996; Robinson and Smyth 2007; Robinson and Smyth 2008) implemented in edgeR using a FDR $< 0.05$ and LogFC$< -0.25$ threshold.

**Pathway enrichment analysis**

Functional enrichment gene-set analysis for GO (Gene Ontology) terms was performed using ToppGene Suite (https://toppgene.cchmc.org/enrichment.jsp) using all *Homo sapiens* genes as background. Statistically significant (Bonferroni q-value $< 0.05$) categories within the GO:Biological Process section were extracted and replotted.

**ChIPseq analysis**

For the ChIPseq analysis, trimming of known adapters and low-quality regions of reads was performed using Fastq-mcf. Sequence quality control was assessed using FastQC (http://www.bioinformatics.babraham.ac.uk/projects/fastqc/). Alignment to the hg19 reference genome was performed using Bowtie 2.2.4 (Langmead and Salzberg 2012). Peaks were called using GEM (Guo et al. 2012) for TFs and BCP (Xing et al. 2012) for GLYR1 and H3K36me3 ChIPseq signals. Read counts per peak were generated with featureCounts (Liao et al. 2014) and normalized to account for differences in sequencing depth between samples using upper quartile normalization separately for the ChIP and input sample. Bound regions were determined using empirical Bayes F-tests for a quasi-likelihood negative binomial generalized log-linear model of the count data as implemented in edgeR. Specifically, we tested for a significant (i.e., non-zero at FDR < 5%) log2 fold-increase in normalized peak signal for ChIP versus the corresponding input sample. 2 or 3 separate samples (and relative inputs) were ran from independent ChIP assays.

- **GATA4 and GLYR1 ChIPseq Genomic Features (Figure S7E)**

We obtained the genomic features associated with GATA4 and GLYR1 ChIP-seq peaks using the annotatr (Cavalcante and Sartor 2017) package in R.

- **Metagene plot analysis (Figure S7D&J and 6D)**

To generate metagene plots, BED files were generated containing regions of interest. The *computeMatrix scale-regions* module of deepTools (Ramírez et al. 2016), which shrinks or stretches all regions in the input BED file to the same length was used to summarize the ChIP signal profile for each region. The ChIP signal was defined in terms of input subtracted tag densities. Specifically, the human genome, hg19 is divided into 20-bp bins. The tag density or normalized difference between the number of the reads in the ChIP sample and the input sample is computed as:

$$\#BinsInGenome \times \left( \frac{\#chipReads}{\#totalChipReads} - \frac{\#inputReads}{\#totalInputReads} \right)$$

- **Analysis of Differential ChIPseq signal (Figure 7A)**

The counts of reads mapping to genes for each of the replicates for each of the ChIPs (Glyr1 and H3K36me3) at the hiPSC and CPC stages were obtained using featureCounts (Liao et al. 2014) using their corresponding aligned reads in bam files. The counts of reads for each of replicates used for assaying gene expression at the two stages in the GSE137920 (Lau et al. 2019) data set were downloaded from GEO (Barrett et al. 2013). Genes where there were not at least two samples with at least 5 (raw) reads in the Glyr1 ChIPs were filtered out from further analyses. The read counts for the remaining genes corresponding to each of the three signals (Glyr1, H3K36m3 and Gene expression) are separately normalized using *calcNormFactors* in edgeR (Robinson and Oshlack 2010). Genes for which the mean Glyr1 signal in their bodies were significantly changed from CPC stage relative to hiPSC stage were determined the likelihood ratio test implemented in edgeR using FDR < 0.1 threshold. The row-normalized log2 transformed Counts-Per-Million (CPM) of Glyr1 signal for these significantly associated (with changing Glyr1 signal) genes were clustered using *kmeans* with 3 clusters implemented in R (R Core Team 2020). The resulting cluster definitions (using the Glyr1 signal) and order of genes were used to visualize the signals (in row-normalized log2 CPM units) in the H3K36me3 and RNA-seq data.

- **Definition of GATA4 and GLYR1 -bound gene categories (list used in most of the panel Figure 6)**

Glyr1 bound genes in Figure 6B are defined as those genes in clusters 1 and 2 in Figure 6A which displayed enriched binding signal at the CPC stage relative to the hiPSC stage. Genes with Gata4 ChIP peaks from the first intron to Transcription End Site (TES) were defined as Gata4 bound genes.

- **Scatter plot analysis (ChIPseq/RNAseq) (Figure S7 A-C)**

Differential gene expression between the hiPSC (day 0) and the CPC stage (day 7) was determined using quasi-likelihood F-test implemented in the *glmQLFTest* in edgeR (Lun et al. 2016) using the count matrix association with the GSE137920 (Lau et al. 2019) filtered for low counts (at least two samples with at least 5 (raw) reads) genes normalized using the *calcNormFactors*. Up-regulated genes were determined using thresholds of 1.5 for log2 fold-change (log2FC > 1.5) and 0.05 for FDR (FDR < 0.05), while down-regulated genes were determined using thresholds of -1.5 for log2 fold-change (log2FC < -1.5) and 0.05 for FDR. The raw read counts for all replicates of Glyr1 at the two stages for all genes that were part of the differential gene expression analyses above, were normalized using *calcNormFactors* and Reads Per Kilobase of transcript, per Million mapped reads (RPKM) was calculated using the *rpkm* function along with the gene lengths based on the Ensembl gene annotation. Similarly, RPKM values were estimated for all replicates of the H3K36me3 ChIP at the two stages. The scatter plots in Figure S7 use the mean log2-transformed RPKM values across replicates at a given stage.

- **Statistical Analysis of GATA4 & GLYR1 ChIPseq overlap (Figure 6B)**

The significance of the overlap of genes bound within their bodies (first intron to Transcription End Site (TES)) by Gata4 and Glyr1 was determined using the Fisher's exact test implemented in R (R Core Team 2020) on 30,611 genes which had detectable reads (5 reads in at least two samples) at the day 7 in the GSE137920 (Lau et al. 2019) data set.

### QUANTIFICATION AND STATISTICAL ANALYSIS

Statistical parameters including the exact value of n, precision measures (mean ± SEM) and statistical significance are reported in the Figures and the Figure Legends. All calculations were performed using R or GraphPad Prism software. When several conditions were to compare, we performed a one-way ANOVA, followed by Tukey range test to assess the significance among pairs of conditions. The significance of the PPIN enrichment in CHD-associated DNVs was calculated with a permutation-based test as explained in the Computational Analysis Methods section. All the p-values related to the violin plots showing features typical of disease genes were obtained using a two-sided Mann-Whitney-Wilcoxon test with Bonferroni correction. The significance of the GATA4 & GLYR1 ChIPseq overlap was estimated using the Fisher.Exact function in R. The level of significance in all graphs is represented as follow: * P<0.05, ** P<0.01, *** P<0.001, **** P<0.0001.

### DATA AND CODE AVAILABILITY

The RNAseq and ChIPseq datasets generated during this study are available at GEO [GSE159411/ https://www.ncbi.nlm.nih.gov/geo/query/acc.cgi?acc=GSE159411]. The mass spectrometry proteomics data have been deposited to the ProteomeXchange Consortium via the PRIDE (Perez-Riverol et al. 2019) partner repository with the dataset identifier PXD022091. Code is available at https://github.com/mepittman/ctf-apms.
